# Thermoprotection by a cell membrane-localized metacaspase in a green alga

**DOI:** 10.1101/2023.04.28.538660

**Authors:** Yong Zou, Igor Sabljić, Natalia Horbach, Adrian N. Dauphinee, Anna Åsman, Lucia Sancho Temino, Marcin Drag, Simon Stael, Marcin Poreba, Jerry Ståhlberg, Peter V. Bozhkov

## Abstract

Caspases are restricted to animals, while other organisms, including plants possess metacaspases (MCAs), a more ancient and broader class of structurally-related yet biochemically distinct proteases. Our current understanding of plant MCAs is derived from studies in streptophytes, and mostly in Arabidopsis expressing nine MCAs with partly redundant activities. In contrast to streptophytes, most chlorophytes contain only one or two hitherto uncharacterized MCAs, providing an excellent platform for MCA research. Here we investigate CrMCA-II, a single type II MCA from a model chlorophyte *Chlamydomonas reinhardtii*. Surprisingly, unlike other studied MCAs and similar to caspases, CrMCA-II dimerizes both *in vitro* and *in vivo*. Furthermore, activation of CrMCA-II *in vivo* correlates with the dimerization. Most of CrMCA-II in the cell is present as a zymogen attached to the plasma membrane (PM). Deletion of *CrMCA-II* by CRISPR/Cas9 compromises thermotolerance leading to increased cell death under heat stress. Adding back either wild-type or catalytically dead CrMCA-II restores thermoprotection, suggesting that its proteolytic activity is dispensable for this effect. Finally, we link the non-proteolytic role of CrMCA-II in thermotolerance to the ability to modulate PM fluidity. Our study reveals an ancient, MCA-dependent thermotolerance mechanism retained by Chlamydomonas and probably lost during the evolution of multicellularity.

## Introduction

In 1992 the first caspase was discovered, interleukin-1*β*-converting enzyme (ICE), better known as caspase-1 (Cerretti et al., 1992; Thornberry et al., 1992), and shortly thereafter, the *Caenorhabditis elegans* cell death gene *ced-3* (Yuan et al., 1993) was identified, heralding a flurry of research on caspases that continues to this day (Julien and Wells, 2017; Van Opdenbosch and Lamkanfi, 2019; Kesavardhana et al., 2020; Ross et al., 2022). However, caspases are a relatively small and mostly animal-specific part of a large superfamily of cysteine proteases (C14 within the CD clan), all sharing a caspase-like structural fold and found throughout all kingdoms of life (Uren et al., 2000; Minina et al., 2020). Among them, eukaryotic metacaspases (MCAs) and prokaryotic MCA-like proteases form the phylogenetically broadest group, with members found in all living organisms except animals (Tsiatsiani et al., 2011; McLuskey and Mottram, 2015; Minina et al., 2017; Klemenčič and Funk, 2019). Despite structural similarity between MCAs and caspases, the two groups of proteases fundamentally differ in their substrate specificity, with MCAs, unlike Asp-specific caspases, cleaving exclusively after positively charged (Arg or Lys) residues (Vercammen et al., 2004; Sundström et al., 2009; Minina et al., 2020).

Whereas all known eukaryotic MCAs contain both p20-(catalytic) and p10-like conserved regions, they are classified into three major types based on the presence of additional protein modules and the relative position of the p20 and p10 regions (Minina et al., 2020). Thus, the distinguishing feature of type I MCAs is the N-terminal pro-domain. Type II MCAs are distinguished by the presence of a long linker separating p20 and p10 regions, whereas type III MCAs are defined by swapping of the two regions, so that the p10 is located N-terminally to the catalytic p20 region. Notably, there is a significant variation in the amino acid sequence, length of distinct protein regions and occurrence of additional motifs among MCAs of the same type, in particular for type I that is presumably responsible for the differences in their biochemical properties and physiological functions (Klemenčič and Funk, 2019). However, further structure-function categorization of the members within each MCA type is not possible today due to a limited success in structural analysis of these enzymes, with only three type I (*Trypanosoma brucei* TbMCA-Ib/MCA2, *Saccharomyces cerevisiae* ScMCA-I/MCA1/YCA1 and *Candida glabrata* CgMCA-I) and two type II (*Arabidopsis thaliana* AtMCA-IIa/MC4 and AtMCA-IIf/MC9) MCA crystal structures available (McLuskey et al., 2012; Wong et al., 2012; Zhu et al., 2020; Conchou et al., 2022; Stael et al., 2023).

Among the three types of MCAs, type I is found in all non-animal organisms, while type II is specific for Chloroplastida (originally named Viridiplantae, Latin for “green plants”) and type III for Heterokontophyta (phytoplanktonic protists; (Choi and Berges, 2013; Minina et al., 2017; Klemenčič and Funk, 2019). Thus, the vast majority of annotated green plant genomes encode both type I and type II MCAs. Green plants form a monophyletic taxon comprising both land plants and green algae re-distributed into two evolutionary lineages, the Chlorophyta (chlorophytes) and Streptophyta (streptophytes), that diverged over a billion years ago (Bemer, 1985; Morris et al., 2018). While the chlorophytes comprise only green algae, the streptophytes include both land plants and the remaining green algae (called “streptophyte algae”; Becker and Marin, 2009).

Research on plant MCAs has so far almost exclusively concerned streptophytes and was dominated by the use of Arabidopsis, due to its advantages of being a genetically tractable model system and having a relatively well-understood biology. The Arabidopsis MCA family consists of nine genes, three of type I (AtMC1-AtMC3 or AtMCA-Ia - AtMCA-Ic, according to original or updated nomenclature, respectively) and six of type II (AtMC4 - AtMC9 or AtMCA-IIa - AtMCA-IIf), the latter including four tandem duplicated genes (AtMC4 - AtMC7 or AtMCA-IIa - AtMCA-IId; (Vercammen et al., 2004; Minina et al., 2020). Arabidopsis MCAs are involved in the regulation of immune and stress responses, aging and programmed cell death (Coll et al., 2010; Watanabe and Lam, 2011; Bollhöner et al., 2013; Coll et al., 2014; Hander et al., 2019; Shen et al., 2019; Wang et al., 2021; Li et al., 2022). For recent reviews on Arabidopsis MCAs see Minina et al., (2017), Salguero-Linares and Coll (2019) and Huh (2022). The absence of strong developmental or fitness phenotypes of single *MCA* gene knockouts in Arabidopsis points to some degree of redundancy among different MCAs (Tsiatsiani et al., 2011; Huh, 2022). Furthermore, while some of the MCA-dependent proteolytic functions might be isoform-specific, it is not easy to prove that experimentally. Indeed, MCAs have rather loose substrate cleavage specificity for residues in P2, P3, and P4 positions (Vercammen et al., 2006; Sundström et al., 2009; Tsiatsiani et al., 2013). This complicates ascribing a given physiological response or developmental event to the proteolytic cleavage of a target protein substrate by a particular MCA, since there is a mixture of MCAs with similar substrate specificity in a cell (or cell lysate).

With these arguments in mind, we turned our attention to the chlorophyte lineage of green plants whose sequenced genomes reveal mostly one or two *MCA* genes (*vs* several in streptophytes; Tsiatsiani et al., 2011), providing a powerful paradigm for plant MCA research. Despite this obvious advantage, there is to date no genetic evidence for the involvement of MCAs in chlorophyte biology, thus precluding evolutionary insight into the MCA-dependent functions in green plants.

We set out to establish chemical and genetic toolboxes for exploring MCAs in a chlorophyte, *Chlamydomonas reinhardtii* (hereafter Chlamydomonas). Chlamydomonas is a unicellular green alga used as a model organism by virtue of having a greatly reduced number of cell types and simplified cell-to-cell communications, as compared to e.g. Arabidopsis (Gutman and Niyogi, 2004), and combining key animal (e.g. cell motility) and plant (e.g. photosynthesis) characteristics (Merchant et al., 2007). The genome of Chlamydomonas encodes one type I and one type II MCA named according to the current nomenclature (Minina et al., 2020), CrMCA-I and CrMCA-II, respectively. Such genetic simplicity renders Chlamydomonas a favourable model for uncovering primordial roles of the two types of MCAs. Here we report biochemical and functional studies of CrMCA-II. Unlike other MCAs, CrMCA-II can oligomerize both *in vitro* and *in vivo*. It is predominantly present as a zymogen, associated with plasma membrane (PM), but translocates to the cytoplasm upon heat stress (HS). Genetic experiments combined with live-cell imaging revealed that CrMCA-II plays a protease-independent cytoprotective role under HS, presumably related to its ability to module PM fluidity.

## Results

### Phylogenetic analysis places two Chlamydomonas MCAs at the base of the evolutionary tree

BLAST search revealed that species in the order Chlamydomonadales, including Chlamydomonas, *Volvox carteri* and *Gonium pectorale*, contain one type I and one type II MCA. Some green algae, e.g. *Raphidocelis subcapitata* and *Auxenochlorella protothecoides*, are completely devoid of type II and contain only type I MCAs. To gain insight into the evolutionary position of Chlamydomonas MCAs, we performed a phylogenetic analysis of MCA sequences from green plants with well-annotated genomes. The analysis demonstrated that both Chlamydomonas MCAs, CrMCA-I and CrMCA-II, as well as other chlorophyte MCAs are situated at the bases of evolutionary clusters comprising type I and type II MCAs from green plants (Supplemental Figure S1).

### CrMCA-II is a Ca^2+^- and redox-dependent arginine-specific protease

We purified bacterially expressed recombinant His-tagged CrMCA-II (rCrMCA-II) using a HisTrap column followed by size exclusion chromatography (SEC). For reasons discussed later, the optimal buffer used for CrMCA-II proteolytic activity measurements and as a basis for further studies was 50 mM Tris pH 7.5, 25 mM NaCl, 20 mM CaCl_2_, 0.1% CHAPS and 7.5 mM DTT. Similar to all hitherto studied MCAs, rCrMCA-II was unable to cleave after Asp (Figure 1A). However, unlike other MCAs, rCrMCA-II displayed a strict preference for Arg over Lys at the P1 position in the tetrapeptide substrates, with only trace hydrolytic activity against Lys-containing substrates (Figure 1A).

**Figure 1.**
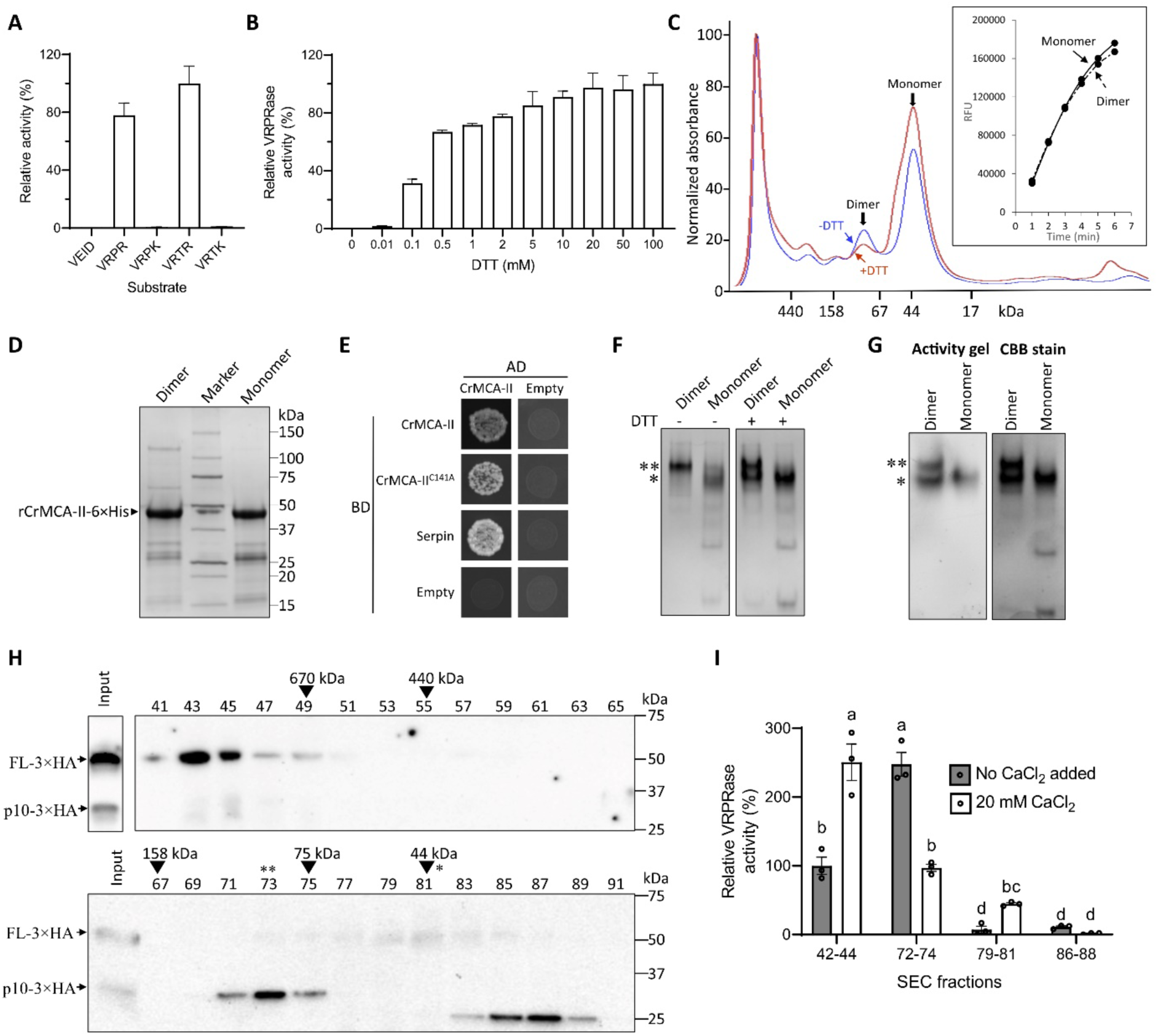
CrMCA-II is a redox-dependent arginine-specific protease prone to oligomerization. **A.** Proteolytic activity of rCrMCA-II against AMC-conjugated tetrapeptide substrates with Arg, Lys, or Asp at the P1 position, relative to Ac-VRTR-AMC, under optimal buffer conditions (50 mM Tris pH 7.5, 25 mM NaCl, 20 mM CaCl_2_, 0.1% CHAPS and 7.5 mM DTT). **B.** Proteolytic activity of rCrMCA-II, as affected by DTT concentration, relative to 100 mM DTT, under optimal buffer conditions, with 50 µM Ac-VRPR-AMC as a substrate. **C.** The size exclusion chromatography (SEC) of rCrMCA-II in the presence and absence of 1 mM DTT. The insert shows proteolytic activity (relative fluorescence units, RFU) of monomeric and dimeric forms of rCrCMA-II separated in the absence of DTT against Ac-VRPR-AMC under optimal buffer conditions. **D.** SDS-PAGE analysis of dimer- and monomer-containing fractions from **C** separated without DTT. **E.** The Y2H assay of CrMCA-II self-interaction and interaction between wild type protease and its catalytically inactive mutant CrMCA-II^C141A^. Interaction between CrMCA-II and serpin, an *in vivo* inhibitor of plant metacaspases, was used as a positive control. **F.** Native PAGE of monomeric and dimeric forms of rCrMCA-II with (+) or without (-) DTT (7.5 mM). **G.** An in-gel proteolytic activity assay of monomeric and dimeric forms of CrMCA-II using native PAGE under optimal buffer conditions, with 50 µM Ac-VRPR-AMC as a substrate. * and ** in **F** and **G** indicate positions of monomer and dimer, respectively. CBB, Coomassie brilliant blue. **H.** Immunoblot analysis of total protein extracts isolated from Chlamydomonas *crmca-ii* mutant strain expressing CrMCA-II-3*×*HA (see Fig. 3) and fractionated by SEC. Uneven fractions from 41 to 91 were separated by SDS-PAGE and analyzed with **α**-HA. The fractions containing protein standards with indicated molecular masses are marked with arrowheads. Based on the calibration curve, the predicted average molecular masses of proteins in fractions 43, 73 and 87 are 1185 kDa, 88.5 kDa and 26.4 kDa, respectively. The theoretical masses of CrMCA-II-3*×*HA zymogen (full-length monomer, FL-3*×*HA) and C-terminal p10 fragment (p10-3*×*HA; generated via auto-cleavage after Arg190) are 46.1 kDa and 25.9 kDa, respectively. The fractions enriched for CrMCA-II-3*×*HA monomer and dimer are denoted by * and **, respectively. **I.** The peptide (Ac-VRPR-AMC) cleavage assay of a subset of SEC fractions from **H** and enriched for megadalton CrMCA-II-3*×*HA assemblies (fractions 42-44), dimer (fractions 72-74), monomer (fractions 79-81), and a fragment smaller than p10 (fractions 86-88). The assay was performed under optimal buffer conditions, with or without addition of 20 mM CaCl_2_. Note the presence of a basal level of cell-derived Ca^2+^ in the assay. Data in **A, B,** and **I** represent the means ± SEM of triplicate measurements. Different letters indicate significant differences at *p* < 0.05 (a two-way ANOVA with Tukey’s honest significant difference test).

As with the majority of MCAs, rCrMCA-II required millimolar concentrations of Ca^2+^ for activation (Supplemental Figure S2A). An inhibitory effect of zinc on CrMCA-II (Supplemental Figure S2B) was not as strong as that for the spruce mcII-Pa (Bozhkov et al., 2005), but similar to human caspase 8 (Stennicke and Salvesan, 1997). rCrMCA-II was active at neutral pH, with an optimum at pH 7.5 (Supplemental Figure S2C), and as expected, was highly sensitive to arginal protease inhibitors (Supplemental Figure S2D).

Noteworthy, rCrMCA-II critically required reducing agents for activation (Supplemental Figure S2E), exhibiting dose-dependent activity with increasing concentration of DTT up to 20 mM (Figure 1B). The predicted structure of the CrMCA-II catalytic center revealed Cys329 in the vicinity of catalytic Cys141 (5.4 Å, Supplemental Figure S3), at a distance even closer than the distance between the catalytic dyad of Cys141 and His87 (6.5 Å), pointing to a possibility of a Cys-Cys bridge formation that could inhibit catalytic activity. However, the rCrMCA-II^C329A^ mutant was significantly less proteolytically active than the WT enzyme and still required reducing conditions for the activity (Supplemental Figure S2F). In agreement with the results obtained for rCrMCA-II, addition of DTT to Chlamydomonas cell lysates prepared from the control (UVM4) strain increased Arg-specific proteolytic activity and hydrogen peroxide had an opposing effect (Supplemental Figure S4). Collectively, these results demonstrate that CrMCA-II is a Ca^2+^- and redox-dependent arginine-specific protease.

### CrMCA-II oligomerizes *in vitro* and *in vivo*

Interestingly, SEC profile of rCrMCA-II in the buffer without DTT and Ca^2+^ displayed two peaks, corresponding to the monomeric (43.5 kDa) and dimeric (87 kDa) forms (Figure 1C). When subjected to SDS-PAGE, both monomer- and dimer-containing SEC fractions displayed full-length protein and autocleavage fragments of a similar molecular mass (Figure 1D). The dimerization was confirmed by Yeast Two-Hybrid (Y2H) assay (Figure 1E). Caspases dimerize through the sixth β-sheet located in the p10 region (MacKenzie and Clark, 2012). We attempted to identify the interaction region of CrMCA-II, but neither p20 nor p10 or linker regions could interact with the full-length CrMCA-II in the Y2H assay (Supplemental Figure S5), indicating that either the corresponding protein fragments are misfolded or the presence of two or even all three regions is required for the dimerization.

The SEC fractions containing either monomeric or dimeric forms of rCrMCA-II were proteolytically active in the buffer supplemented with DTT and Ca^2+^ (Figure 1C, insert). As DTT may prevent protein-protein interaction by disrupting intermolecular disulfide bonds, the activity of the dimer-containing fraction may in fact be caused by monomers forming upon addition of DTT. Indeed, we observed a decreased dimer-to-monomer ratio when 1mM DTT was added to the SEC buffer, confirming that DTT induces dimer-to-monomer transition (Figure 1C). This was further confirmed using native gel electrophoresis (Figure 1F). To unequivocally determine which of the two forms of rCrCMA-II represent catalytically active protease, we performed an in-gel activity assay wherein monomeric and dimeric forms of rCrMCA-II were subjected to native electrophoresis followed by the addition of Ac-VRPR-AMC. Both monomers and dimers were able to cleave the fluorescent substrate (Figure 1G), demonstrating that dimerization is dispensable for rCrMCA-II activation *in vitro*.

To investigate whether CrMCA-II dimerizes *in vivo*, we performed SEC of total protein extract from a knockout Chlamydomonas strain complemented with HA-tagged WT CrMCA-II (CrMCA-II-3*×*HA; generation of transgenic strains is discussed later), followed by SDS-PAGE and immunoblot analysis of the SEC fractions with anti-HA (Figure 1H). The analysis identified three major forms of CrMCA-II-3*×*HA co-existing in the cell extract: monomer (peak fraction 81), dimer (peak fraction 73) and a megadalton-scale complex (peak fraction 43; Figure 1H). While CrMCA-II-3*×*HA in the SEC fractions containing monomer and megadalton complex was present mainly as a zymogen (i.e. full-length protein), the dimer-containing fractions were represented by the autoprocessed, mature protease, generating a p10-fragment under denatured conditions that is recognized by *α*-HA (Figure 1H). This fragment results from the autocleavage at a conserved site (after Arg190 in case of CrMCA-II and based on sequence alignment with known cleavage sites from Arabidopsis type II MCAs) within the linker region of type II MCAs (Zhu et al., 2020; Supplemental Figure S6. Besides three major forms of CrMCA-II-3*×*HA, the SEC analysis of the total protein extract has also revealed an abundance of a C-terminal fragment smaller than the p10-fragment present in a free, unbound form (peak fraction 87; Figure 1H) and apparently representing a product of self-degradation (Watanabe and Lam, 2011).

Since endogenous CrMCA-II exists as monomeric, dimeric and large (megadalton-scale) species, we asked whether these species are proteolytically active in the substrate cleavage assay, at two contrasting levels of Ca^2+^, but otherwise maintaining optimal buffer conditions: (i) without adding any exogenous Ca^2+^, and (ii) with addition of 20 mM CaCl_2_. While the first conditions approximate the chemical environment *in situ*, with only a basal level of cell-derived Ca^2+^ present in the assay, the second conditions provide optimal level of Ca^2+^ for detecting full activity of a proteolytically competent CrMCA-II. Ca^2+^ supplementation strongly enhanced VRPRase activity of the SEC fractions with monomeric and megadalton CrMCA-II species, largely represented by zymogen, but inhibited activity of dimer-containing fractions represented by the autoprocessed mature enzyme (Figure 1I). As expected, the SEC fractions containing a C-terminal fragment of CrMCA-II smaller than p10 displayed only trace activity, independently of the Ca^2+^ level (Figure 1I).

Results of SEC, immunoblotting and proteolytic activity assays together have two important implications. First, the pattern of the CrMCA-II protein in SDS-PAGE can be used as a proxy to its proteolytic competence. Second, it is highly likely that it is a dimer representing a proteolytically active form of CrMCA-II *in vivo*, whereas monomeric and megadalton CrMCA-II species are present as catalytically competent but inactive zymogen molecules.

### Development of CrMCA-II specific chemical probes

To determine substrate specificity of CrMCA-II beyond P1 position and to develop optimal substrates, hybrid combinatorial substrate library (HyCoSuL) screening combining both natural and unnatural amino acids, was employed (Poreba et al., 2017). Using the library with fixed Arg at the P1 position, the enzyme preferences at the P2, P3 and P4 positions were screened (Supplemental Figure S7). A broad substrate specificity was found for rCrMCA-II from P4 to P2 positions. Nevertheless, to get a better insight into enzyme preferences, a set of individual tetrapeptide substrates bearing a 7-Amino-4-carbamoylmethylcoumarin (ACC) fluorophore reporter were synthesized and subjected to rCrMCA-II hydrolysis (Figure 2A-C; Supplemental Figure S8; Supplemental Table S1). This screening enabled to identify two substrates, one composed of natural amino acids, Ac-His-Arg-Thr-Arg-ACC (Ac-HRTR-ACC), and one with unnatural amino acids, Ac-His(Bzl)-hSer(Bzl)-Thr-Arg-ACC (Ac-H(Bzl)-hS(Bzl)-TR-ACC), both of which show a high cleavage rate by rCrMCA-II. The catalytic activity of rCrMCA-II on these substrates was significantly higher than that on the commercially available and commonly used MCA and paracaspase substrate Ac-VRPR-ACC (Table 1, Figure 2B, C), which was designed from the optimal substrate preference of Arabidopsis AtMCA-IIf/MC9 (Vercammen et al., 2006), making them efficient probes for measuring CrMCA-II activity.

**Figure 2.**
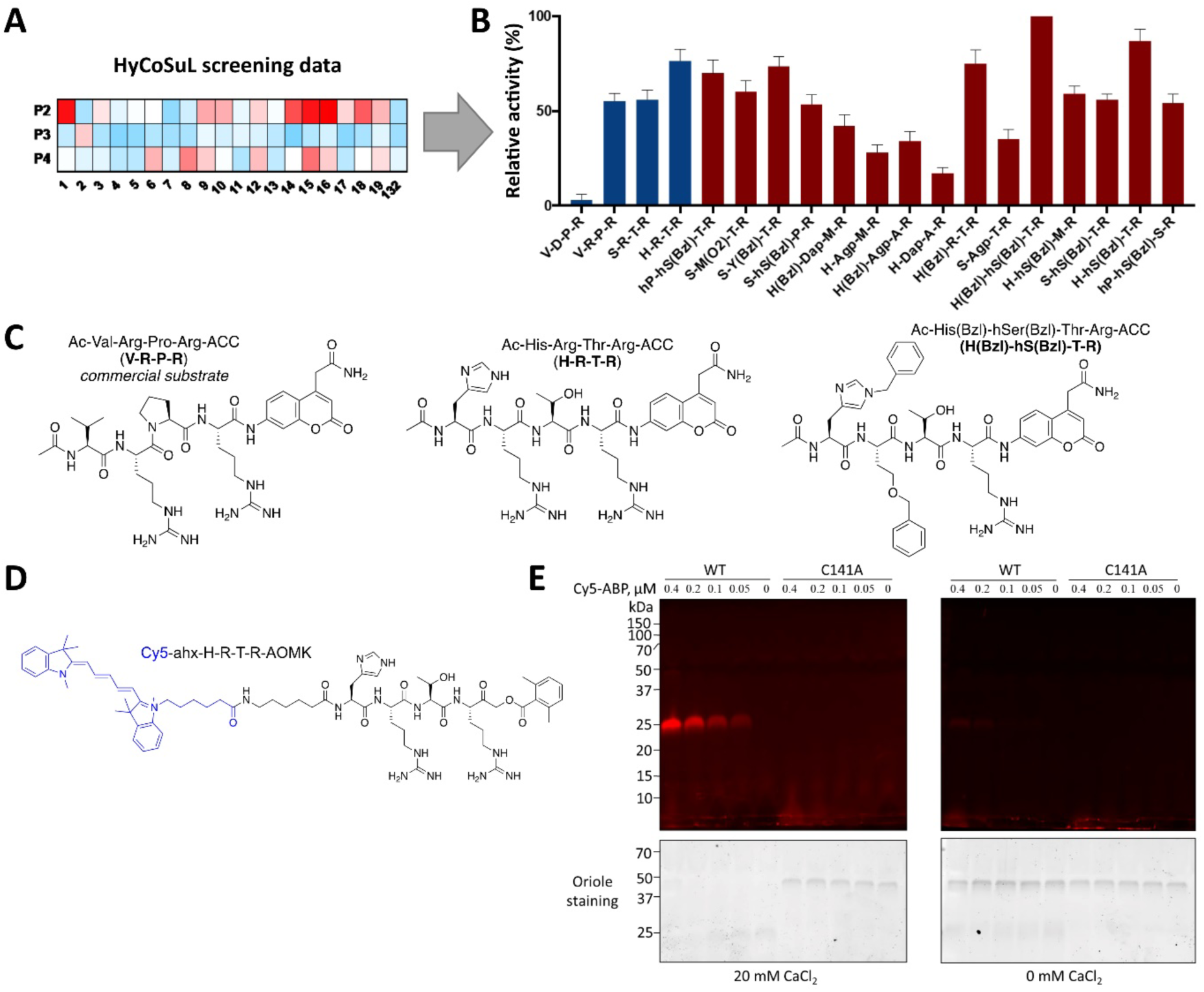
A chemical toolbox for CrMCA-II. **A.** HyCoSuL screening data were used to extract optimal peptide sequences for CrMCA-II substrates and activity-based probes. **B.** Proteolytic activity of rCrMCA-II against individual fluorescent substrates containing natural (blue bars) and unnatural amino acids (red bars). Data represent the means ± SEM of triplicate measurements. **C.** The structures of rCrMCA-II substrates: Ac-VRPR-ACC (commercially available as Ac-VRPR-AMC), Ac-HRTR-ACC (most preferred with natural amino acids), and Ac-H(Bzl)-hS(Bzl)-TR-ACC (most preferred with unnatural amino acids). **D.** The structure of Cy5-labelled covalent activity-based probe (Cy5-ABP) with HRTR peptide motif and AOMK electrophilic warhead. **E.** The binding of Cy5-ABP at different concentrations to 0.4 µM wild-type (WT) rCrMCA-II or its catalytically inactive mutant (C141A) under optimal buffer conditions, in the absence or presence of 20 mM CaCl_2_, visualized after SDS-PAGE.

**Table 1.**
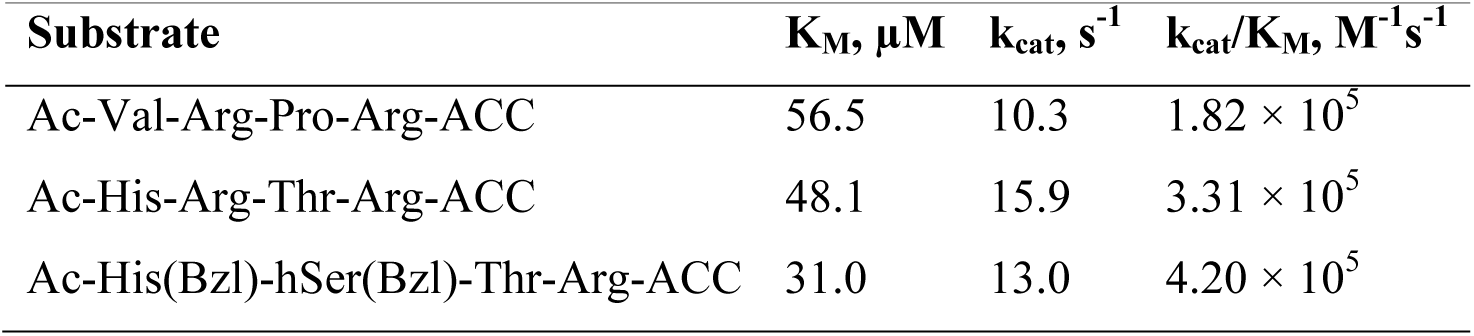
The catalytic activity of rCrMCA-II against three selected substrates.

A considerable benefit of analyzing protease substrate preferences using HyCoSuL technology is the development of a peptide that perfectly fits to the active site of the investigated enzymes. Therefore, the rCrMCA-II preferred peptidic substrates were further used to design fluorescently labeled activity-based probes (ABPs), *viz.* covalent inhibitors that bind to the enzyme active site in an irreversible manner. To do this, we labeled HRTR and H(Bzl)-hS(Bzl)-TR peptides with a cyanine 5 (Cy5) tag through an 6-aminohexanoic acid (Ahx) linker to separate the fluorophore and peptide. We selected the acyloxymethyl ketone group (AOMK) to use as the warhead (Figure 2D), as this electrophile is known to covalently react with the catalytic Cys residue of Cys proteases (Kato et al., 2005). In such a manner, we developed two CrMCA-II ABPs, Cy5-ahx-HRTR-AOMK and Cy5-ahx-H(Bzl)-hS(Bzl)-TR-AOMK (Figure 2D and Supplemental Figure S9.

To determine whether the ABPs can differentiate between WT and catalytically dead mutant of CrMCA-II, as well as between mature WT CrMCA-II and its zymogen, we incubated purified rCrMCA-II or rCrMCA-II^C141A^ with Cy5-ahx-HRTR-AOMK under optimal buffer conditions in the presence or absence of Ca^2+^ followed by SDS-PAGE. As shown in Figure 2E, no labelling was observed for rCrMCA-II^C141A^ with or without Ca^2+^, whereas the ABP labelled only the autoprocessed, mature form of rCrMCA-II generated in the presence of Ca^2+^ in a dose dependent manner. This indicates that the ABP can distinguish between active rCrMCA-II and zymogen. The fragment of mature rCrMCA-II covalently bound by the Cy5 probe is expected to include the p20 region as well as a part of the linker to Arg190, migrating as a band of ∼ 25 kDa (Figure 2E). These results demonstrate that our Cy5-labeled ABPs are useful tools for detecting mature CrMCA-II enzyme.

### Generation of *CrMCA-II* knockout (KO) mutants and complementation strains

To study physiological roles of CrMCA-II, we generated *crmca-ii* mutant strains using the recently established targeted insertional mutagenesis method based on the CRISPR/Cas9 system (Picariello et al., 2020). We obtained three strains (*ii-4*, *ii-9* and *ii-23*) lacking *CrMCA-II* expression. PCR amplification of genomic DNA from strains *ii-9* and *ii-23* with a pair of primers flanking the insertion site furnished a product of around 3.5 kbp (*vs.* 1.8 kbp in the control strain, Figure 3A, B). Sequencing confirmed the insertion of AphVIII cassette conferring paromomycin resistance in the genomic DNA (Supplemental Figure S10). A third mutant, *ii-4* showed no product after colony PCR (Figure 3B), indicating a larger insertion.

**Figure 3.**
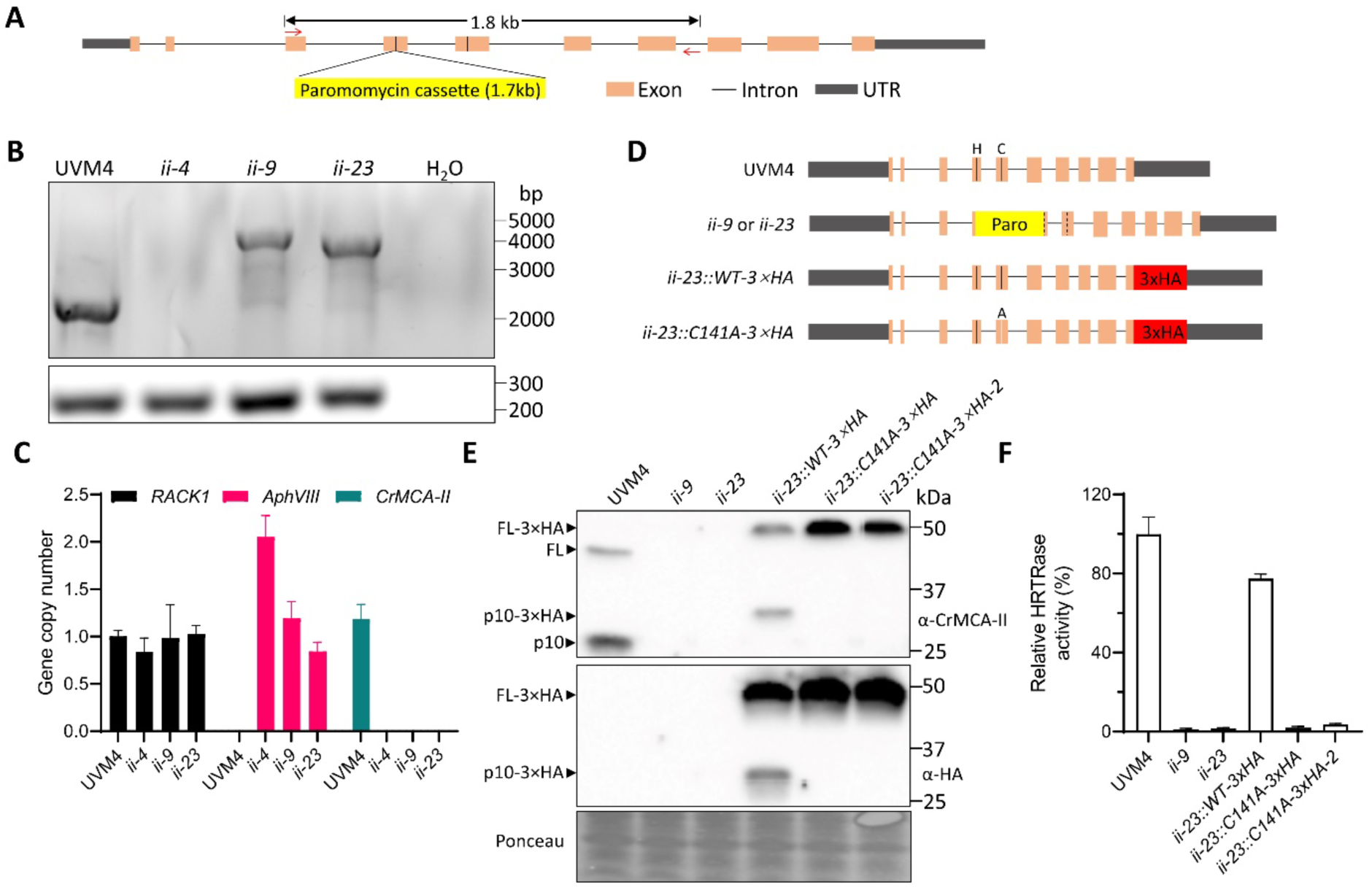
The generation of *crmca-ii* mutant and complementation strains in UVM4 background. **A.** Schematic illustration of the editing site of *CrMCA-II* gene and a pair of primers (red arrows) used for PCR analysis. The vertical lines in exons 4 and 5 indicate the position of a catalytic dyad of His87 and Cys141. **B.** PCR test using the primer pair depicted in **A** with genomic DNA as a template. Distilled water was used as a negative control. Bottom panel shows PCR product of *RACK1* amplification (positive control). *ii-4*, *ii-9* and *ii-23* are presumed mutant strains. **C.** Real-time quantitative PCR (RT-qPCR) analysis of *RACK1*, *AphvIII* and *CrMCA-II* genes using genomic DNA as a template to evaluate genome insertion events. Data represent the means ± SEM of triplicate measurements. Paro, paromomycin resistance cassette. **D.** Schematic illustration of *CrMCA-II* gene structure in the generated strains. The vertical lines indicate position of a catalytic dyad of His87 (H) and Cys141 (C), and a substitution of Cys141 for Ala (A) in strains complemented with a catalytically inactive mutant. The dashed vertical lines corresponding to a catalytic dyad indicate abolished transcription in the mutant strains. **E.** Immunoblot analysis of CrMCA-II in UVM4, *CrMCA-II* mutant (*ii-9* and *ii-23*), and complementation strains (one *ii-23::WT-3×HA* strain and two *ii-23::C141A-3×HA* strains) using **α**-CrMCA-II and **α**-HA. Ponceau staining was used as a loading control. **F.** The peptide (Ac-HRTR-ACC) cleavage assay of cell lysates (total protein extracts) from the same strains as in **E.** Data represent the means ± SEM from three independent biological replicates.

Quantitative PCR (qPCR) has been adapted for estimating gene copy number in a genome, as an alternative to a more time-consuming Southern blot (Hoebeeck et al., 2007). Accordingly, we used this method for assessing insertion events during genome editing. While *ii-9* and *ii-23* strains contained a single *AphVIII* gene copy, confirming single insertion events, *ii-4* presumably had two *AphVIII* gene copies inserted into the genome (Figure 3C, D). Furthermore, qPCR analysis confirmed that none of the three *crmca-ii* mutant strains contained an intact *CrMCA-II* gene (Figure 3C).

To verify a lack of the CrMCA-II protein in the generated mutants, we used an antibody raised against a 16 amino acid-long peptide from the p10 region of CrMCA-II (Supplemental Figure S11A). The antibody recognized both the zymogen (∼ 42.4 kDa) and a fragment composed of p10 and a part of the linker region generated *via* autocleavage (migrated as ∼ 25 kDa but estimated to be 22 kDa) on immunoblots of total protein extracts isolated from the control strain (UVM4; Figure 3E and Supplemental Figure S11B). In contrast, none of the three *crmca-ii* strains revealed the presence of CrMCA-II protein (Figure 3E and Supplemental Figure S11B), indicating that all three strains were KO mutants. We also analyzed cell lysates isolated from the control strain and KO mutants for proteolytic activity of CrMCA-II using the optimized substrate Ac-HRTR-ACC. To minimize risk that cleavage of this substrate by other Chlamydomonas proteases could mask the effect of CrMCA-II deficiency, we blocked activity of aspartate, cysteine and serine proteases, except for arginal proteases, using a combination of aprotinin, pepstatin A, PMSF and Pefabloc SC. While HRTRase activity was readily detected in the lysates from UVM4 strain, it was nearly absent in the lysates from the *crmca-ii* strains (Figure 3F and Supplemental Figure S11C), making them a robust tool for functional studies.

To further expand our toolbox, we cloned the native *CrMCA-II* promoter and genomic DNA of *CrMCA-II* into MoClo system (Crozet et al., 2018), and obtained mutant strains complemented with HA-tagged WT CrMCA-II or catalytically dead CrMCA-II^C141A^ (Figure 3D). While complementation with WT enzyme furnished mature, catalytically active protease, complementation with autoprocessing-deficient CrMCA-II^C141A^ could not restore HRTRase activity in cell lysates (Figure 3E, F).

### CrMCA-II plays cytoprotective role under heat stress (HS) independent of its protease activity

We took advantage of the generated probes and algal strains to explore the role of CrMCA-II *in vivo*. To begin, we compared cell growth of UVM4 and mutant strains under normal conditions (TAP medium, pH 7.0, 23°C) and different types of abiotic stress, including salt stress, nitrogen, phosphorous or acetate starvation, high (pH 8.4) or low (pH 6.0) pH stress, and HS (42°C). We only observed a significant difference in the growth rate under HS, with *crmca-ii* strains exhibiting suppressed growth as compared to UVM4 (data not shown). We then compared the frequency of cell death in UVM4, *crmca-ii* and complemented strains after 60-min HS at 42°C using fluorescein diacetate (FDA) staining. We found that the mutant strains had a higher frequency of cell death than the UVM4 strain (Figure 4A). Consistently, complementation with WT CrMCA-II restored cell viability. Surprisingly, two independent strains complemented with CrMCA-II^C141A^ did so too (Figure 4A), demonstrating that the cytoprotective effect of CrMCA-II under HS conditions is independent of its protease activity.

**Figure 4.**
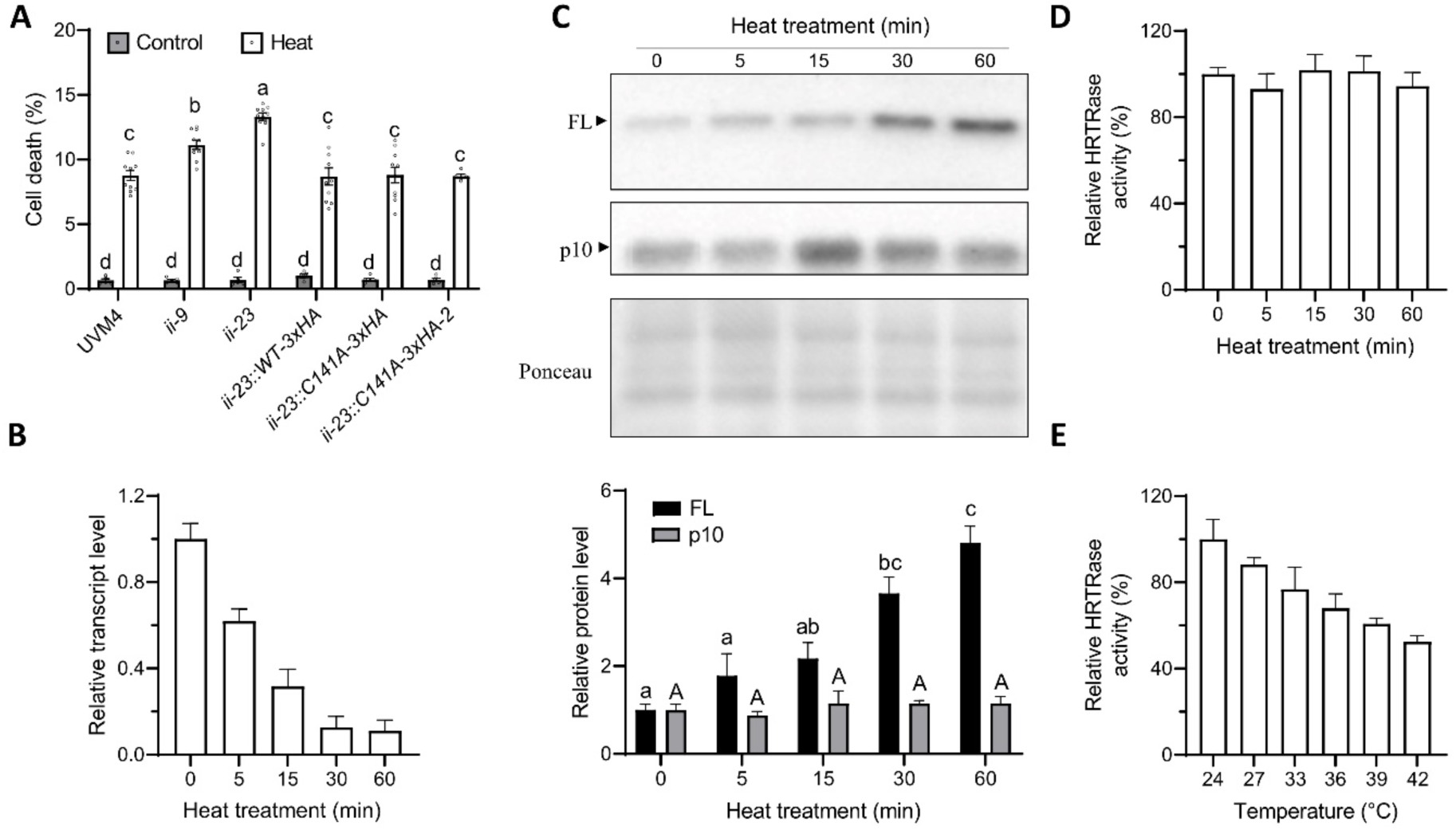
CrMCA-II plays a cytoprotective role during HS. **A.** The frequency of cell death in the UVM4, mutant and complementation strains growing under control conditions (23°C; Control) or subjected to 60-min HS at 42°C (Heat), as determined by FDA staining. **B.** The relative transcript levels of *CrMCA-II* in the UVM4 strain at 0, 5, 15, 30 and 60 min HS at 42°C determined by RT-qPCR. The transcript level at 0 min was set to 1. **C.** Immunoblot analysis of full-length (FL) and p10 fragment of CrMCA-II in total protein extracts isolated from the UVM4 strain at 0, 5, 15, 30, and 60 min of HS at 42 °C. The chart shows relative levels of the FL and p10 based on densitometry analysis of the corresponding bands. The levels of FL and p10 at 0 min were set to 1, respectively. Ponceau staining was used as a loading control. **D.** The relative HRTRase activity of cell lysates isolated from the UVM4 strain at 0, 5, 15, 30, and 60 min of HS at 42°C. **E.** The relative HRTRase activity of rCrMCA-II pre-treated for 5 min at different temperatures in Ca^2+^- and DTT-free buffer (non-permissive conditions) before activity measurements under optimal buffer conditions. The data shown on charts **A** and **C** were analyzed using a two-way ANOVA with Tukey’s honest significant difference test. Different letters indicate significant differences at *p* < 0.05. Data on charts **A** and **C** represent the means ± SEM from three independent biological replicates, with each replicate in case of **A** including triplicate measurements. Different letters indicate significant differences at *p* < 0.05 (a two-way ANOVA with Tukey’s honest significant difference test). Data in **B**, **D** and **E** show the means ± SEM of one representative experiment with triplicate measurements.

Next, we wondered what happened to CrMCA-II during 42°C HS at the transcriptional and protein levels. We found that while HS led to a steady decrease of the *CrMCA-II* transcript level in the UVM4 strain (Figure 4B), it promoted zymogen accumulation without significantly altering the abundance of the mature form of the enzyme, detected through the formation of p10 fragment recognized by *α*-CrMCA-II (Figure 4C; Supplemental Figure S11B). In agreement, HRTRase activity of cell lysates remained constant over the 60-min duration of HS treatment (Figure 4D). These data indicated that HS triggered a *trans*-acting mechanism and/or conformational changes suppressing CrMCA-II activation and leads to zymogen accumulation. To clarify this further, we measured HRTRase activity of purified rCrMCA-II pretreated for 5 min in Ca^2+^- and DTT-free buffer (non-permissive conditions) at various temperatures ranging from 24°C to 42°C and observed a decrease in the activity with increased pre-treatment temperature following addition of Ca^2+^ and DTT (Figure 4E). We thus assumed that heat-induced suppression of CrMCA-II activation could be due to conformational changes of the zymogen.

### CrMCA-II is associated with the plasma membrane (PM) but translocates to the cytoplasm during HS

Given the thermoprotective role of CrMCA-II, we wondered where it was located in a Chlamydomonas cell before and after onset of HS. For studying subcellular localization of CrMCA-II, we assembled an expression vector with a constitutive *P_PSAD_* promoter, gDNA of *CrMCA-II*, mVenus tag and *T_PSAD_* terminator. After transformation into UVM4 cells (Neupert et al., 2009), we obtained a strain overexpressing CrMCA-II with a C-terminal mVenus tag, that we named CrMCA-II-overexpressor 14-3 (OE14-3). Confocal microscopy analysis of unstressed OE14-3 showed a strong mVenus signal co-localized with FM4-64-stained plasma membrane (PM, Figure 5A). PM localization of CrMCA-II-mVenus was confirmed using strain CC-4533, distinguished from UVM4 by the presence of both an intact cell wall and flagella (Supplemental Figure S12), suggesting that this localization is a strain-independent characteristic of Chlamydomonas.

**Figure 5.**
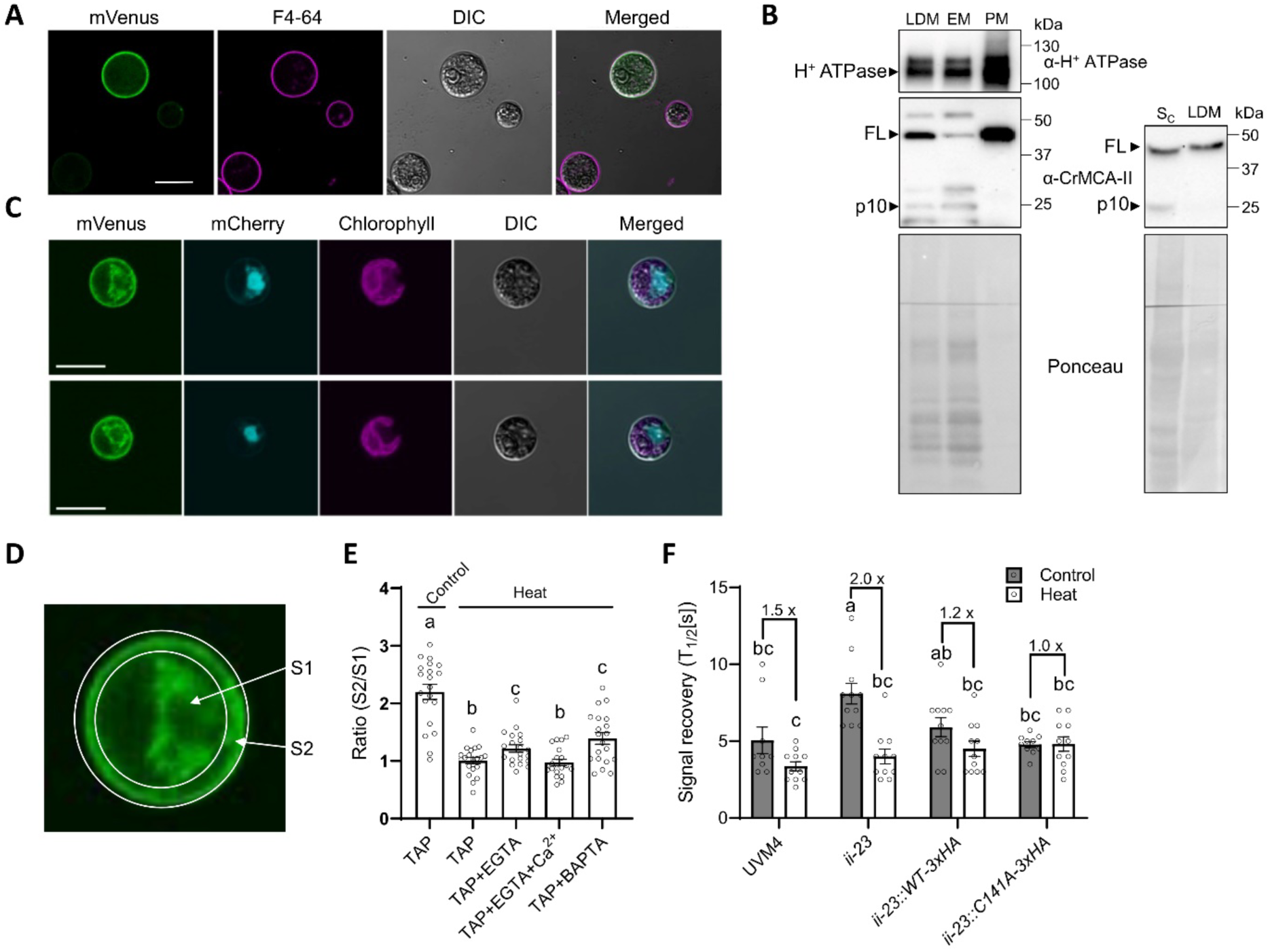
CrMCA-II is localized to the PM and modulates its fluidity. **A.** The PM localization of CrMCA-II-mVenus (green) in unstressed UVM4 cells (grown at 23°C). Cells were stained with FM4-64 (magenta). Scale bars, 10 µm. **B.** Immunoblot analysis of H^+^ ATPase and CrMCA-II in different sub-cellular fractions prepared from unstressed UVM4 cells. LDM, low-density membrane fraction; PM, plasma membrane fraction; EM, endomembrane fraction; S_c_, concentrated soluble proteins. Ponceau staining was used as a loading control. FL, full-length CrMCA-II. **C.** The PM-to-cytoplasm translocation of CrMCA-II-mVenus (green) after HS at 42°C for 60 min. Cells co-expressed mCherry-NLS (turquoise). Scale bars, 10 µm. **D.** Separation of the whole cell area into the cytoplasm area (S1) and PM area (S2) for CrMCA-II-mVenus PM-to-cytoplasm signal ratio quantification in **E. E.** The PM-to-cytoplasm ratio of CrMCA-II-mVenus in unstressed (23°C, Control) and heat-stressed (42°C, Heat) cells grown in TAP medium (0.34 mM CaCl_2_) with or without EGTA (0.75 mM), BAPTA (0.5 mM), or simultaneous presence of both EGTA (0.75 mM) and CaCl_2_ (1 mM). **F.** Recovery rate of fluorescence signal of DiOC6(3) following photo-bleaching (FRAP) of cells from indicated strains grown under no-stress conditions (23°C, Control) or HS (39°C for 60 min, Heat). Numbers on the top of the chart indicate fold-increase of PM fluidity in heat stressed compared to unstressed cells. Data in **E** and **F** represent the means ± SEM from at least three independent biological replicates, each containing at least 3 individual measurements (cells). Different letters indicate significant differences at *p* < 0.05 (a two-way ANOVA with Tukey’s honest significant difference test).

To provide independent biochemical evidence for the PM localization of CrMCA-II, proteins from several fractions including cytosolic, low-density membrane [LDM, including PM and endomembranes (EM)], and separated PM and EM (Supplemental Figure S13) were immunoblotted with antibodies against CrMCA-II and H^+^ ATPase, a marker for PM (Norling et al., 1996). As expected, H^+^ ATPase was enriched in the PM fraction, which also displayed a strong accumulation of CrMCA-II, in agreement with the microscopy data (Figure 5B, left blot). Although confocal microscopy failed to detect cytoplasmic CrMCA-II-mVenus signal in unstressed cells (Figure 5A), the concentrated soluble protein fraction (precipitated and dissolved in the same volume as LDM fraction) exhibited a similar level of CrMCA-II as LDM fraction (Figure 5B, right). Interestingly, CrMCA-II in the PM fraction was present only as a zymogen, in contrast to the cytosolic and EM fractions containing both zymogen and mature enzyme (represented by p10 fragment; Figure 5B). Taken together, these results confirm PM localization of CrMCA-II in unstressed cells, but also point to the existence of the cytoplasmic pool of the protein. In addition, the data suggest that while the PM pool of CrMCA-II is represented exclusively by zymogen, the cytoplasmic pool is partitioned between zymogen and mature protease.

The localization of CrMCA-II under HS was studied using a strain overexpressing mCherry fused to simian virus 40 nuclear localization signal (SV40), in the OE14-3 background. We observed that when mid-log grown cells were exposed to HS at 42°C for 60 min, a strong mVenus signal appeared in the perinuclear cytoplasm (Figure 5C). Thus, the ratio of PM to cytoplasmic fluorescent signals of CrMCA-II was decreased by more than two times in the heat-treated compared to unstressed cells (Figure 5D, E). Importantly, proteolytic activity of CrMCA-II was not required for its PM-to-cytoplasm re-distribution, as revealed by the microscopy analysis of a strain expressing mVenus tagged CrMCA-II^C141A^ (Supplemental Figure S14A-C).

Next, to determine whether the HS-induced accumulation of the cytoplasmic CrMCA-II is a consequence of its translocation from the PM or *de novo* protein synthesis, the cells were treated with cycloheximide (CHX), an inhibitor of translation (Schneider-Poetsch et al., 2010), prior to HS. Since CHX could not prevent or alleviate perinuclear signal (Supplemental Figure S14D, E), we conclude that enhanced cytoplasmic accumulation of CrMCA-II during HS is a result of PM-to-cytoplasm translocation rather than *de novo* protein synthesis.

HS is known to induce the influx of Ca^2+^ through PM (Saidi et al., 2009). MCAs generally contain two Ca^2+^ binding sites, one with high affinity and unknown function and another with low affinity and involved in catalytic activation (Klemenčič and Funk, 2019). During HS, Ca^2+^ influx results in an increase of intracellular Ca^2+^ concentration to a micromolar level (summarized in Schroda et al., 2015), i.e. to the range where MCAs bind Ca^2+^ with high affinity. To test whether the influx of Ca^2+^ through PM is involved in the PM-to-cytoplasm translocation of CrMCA-II, log-phase OE14-3 cells were pre-treated with Ca^2+^ chelators 0.75 mM EGTA or 1,2-bis(o-aminophenoxy) ethane-N,N,N,N′-tetraacetic acid (BAPTA) prior to HS. Calcium chelation led to a moderate but statistically significant suppression of the CrMCA-II translocation, whereas addition of excess Ca^2+^ to the EGTA-treated cells fully reverted that effect (Figure 5E), suggesting that Ca^2+^ influx during HS does play a role in the PM-to-cytoplasm translocation of CrMCA-II.

### CrMCA-II modulates PM fluidity

We hypothesized that the molecular mechanism underlying the cytoprotective effect of CrMCA-II under HS might be related to its potential role in modulating PM fluidity, a dynamic property of PM critically involved in HS signaling (Saidi et al., 2009; Niu and Xiang, 2018) and activation of cell death (Tekpli et al., 2013). To address this hypothesis, we stained the algal strains with DiOC6(3), a lipophilic green fluorescent dye with a high affinity to Chlamydomonas PM (Long et al., 2016), followed by a fluorescence recovery after photo-bleaching (FRAP) assay. An attempt of running the assay at 42°C failed due to hypersensitivity of the heat-treated cells to the PM photobleaching that resulted in cell collapse, so we reduced the HS temperature to 39°C.

We did not observe any significant differences in the proportion of the initial signal recovery of DiOC6(3) on the PM between different strains and regardless of the temperature (data not shown). However, CrMCA-II deficiency led to a strong decrease of signal recovery rate (*viz.* T_1/2_ increase) of DiOC6(3) in unstressed cells, as compared to UVM4 and both WT and CrMCA-II^C141A^ complementation strains (Figure 5F). While UVM4 and complementation strains generally responded to HS by a mild increase in the PM fluidity, this increase was significantly augmented in the CrMCA-II deficient mutant (Figure 5F, Supplemental Figure S15). These results indicate that CrMCA-II controls PM fluidity by preventing PM rigidization in unstressed conditions, which also enables to avoid the hyperfluidization in response to HS. Similar to cytoprotection and PM localization, this function of CrMCA-II occurs independently of its protease activity.

## Discussion

### Oligomerization of CrMCA-II

The absence of oligomerization-dependent activation of MCAs has been proposed as a generic feature distinguishing them from caspases and paracaspases (McLuskey and Mottram, 2015; Minina et al., 2020). Our observations with Chlamydomonas CrMCA-II are not fully consistent with this notion. Firstly, bacterially expressed CrMCA-II can form monomers and dimers, which are both catalytically active (Figure 1C, D, G). While dimerization in this case may be due to a high concentration of the recombinant protein, the results of Y2H assay do not support this assumption (Figure 1E). Secondly, Chlamydomonas protein extracts contain monomeric, dimeric and megadalton-size CrMCA-II species, of which only dimers seem to be fully active (Figure 1H, I).

Our observation that dimerization of endogenous CrMCA-II correlates with the autocleavage yielding the p10 fragment (Figure 1H) is in agreement with a hypothetical model for type II MCA maturation, which postulated that autocleaved MCA molecules could form a catalytically active homodimer (Lam and Zhang, 2012). One cannot completely rule out a possibility that protein species from Chlamydomonas cell extracts with a molecular mass corresponding to CrMCA-II dimer and recognized by specific antibody are in fact heterodimers composed of CrMCA-II and not an unknown protein of a similar molecular mass, although the likelihood of such scenario is low.

The structural details of CrMCA-II dimerization remain unknown. While type II MCAs have a similar β-strand arrangement with caspases, the L2 loop is embedded in the potential dimerization interface, making it impossible for the dimerization to occur in the same way as in caspases (Zhu et al., 2020). However, unlike caspases, type II MCAs have a long, intrinsically disordered linker between the p20 and p10 regions (Supplemental Figure S3), providing an alternative interface for dimerization. Although we failed to detect interaction between the individually expressed linker and the full-length CrMCA-II in the Y2H assay (Supplemental Figure S5), this could be due to large conformational changes of the disordered linker in yeast. Structural data for catalytically mature CrMCA-II are required to unambiguously determine which structural elements serve as the dimerization interface, as well as molecular topology of the dimer. Likewise, structure-function understanding of the megadalton-scale, CrMCA-II zymogen-containing complex identified in Chlamydomonas extracts awaits further studies.

### CrMCA-II is a redox-dependent protease

Reducing conditions are crucial for CrMCA-II activation both *in vitro* (Supplemental Figure S2E) and in cell lysates (Supplemental Figure S4), and at the same time stimulate dimer-to-monomer transition of rCrMCA-II (Figure 1E), suggesting that Cys-Cys disulfide bond(s) might contribute to the CrMCA-II dimerization. The redox-regulated activity of a type III MCA from the marine diatom *Phaeodactylum tricornutum* was previously found to correlate with the formation of a disulfide bond between Cys202 and Cys259 (Graff van Creveld et al., 2021). While these Cys residues are not conserved in CrMCA-II (not shown), we reasoned that the formation of a disulfide bond between catalytic Cys141 and the closely situated Cys329 would explain strict requirement of reducing conditions for the protease activity of CrMCA-II (Supplemental Figure S3). However, Cys329Ala mutation did not render protease activity independent of the reducing agent (Supplemental Figure S2F), calling for further research to uncover an underlying mechanism and its physiological implications *in vivo*.

Noteworthy, strong redox dependence of CrMCA-II proteolytic activity revealed in our study is consistent with the appearance of CrMCA-II in the published redox thiol proteome datasets of Chlamydomonas (Pérez-Pérez et al., 2017; McConnell et al., 2018). Furthermore, redox dependence might be an important regulatory mechanism related to CrMCA-II location on the PM; a major site for ROS production, perception and signal transduction in a cell (Nordzieke and Medraño-Fernandez, 2018), and responsible for thermotolerance and other physiological functions of this MCA yet to be identified.

### Substrate specificity of CrMCA-II

Without regard to the non-proteolytic role in thermotolerance, mature CrMCA-II exhibits a multifold higher preference for Arg than for Lys at the P1 position in tetrapeptide substrates (Figure 1A), implying that the S1 binding pocket of CrMCA-II has a higher affinity for Arg than for Lys (Stael et al., 2023). Whether or not this holds true *in vivo* is unknown and necessitates information about cleavage sites in the native targets. In fact, strong preference for Arg over Lys in peptidic substrates is a common feature of MCAs (Minina et al., 2017), which however, is not obvious when analyzing cleavage sites of the target proteins (Sundström et al., 2009; Tsiatsiani et al., 2013). This suggests that the residues at P1’ to P4’ of the substrates may also affect the binding and activity of MCAs during cleavage.

As for a specificity beyond P1, CrMCA-II displayed a clear preference for a basic amino acid at P3 site (Supplemental Figure S7), similar to Arabidopsis AtMCA-IIf/MC9 (Vercammen et al., 2006). Accordingly, substitution of Arg for Asp at P3 of the commercially available VRPR-based substrate greatly diminished proteolytic activity of CrMCA-II (Figure 2B). However, unlike AtMCA-IIf, which prefers Pro, Tyr or Phe at P2 and branched chain Ile or Val at P4 site (Vercammen et al., 2006), CrMCA-II instead favours substrates with Ala, Thr or Ser at P2 and His or Ser at P4 (Supplemental Figure S7). This comparison implies that while MCAs have rather loose P2-P4 substrate specificity, it differs among different class members.

The ABPs designed from the substrate specificity screening are useful tools for detecting catalytically active proteoforms of the recombinantly produced CrMCA-II (Figure 2D, E). Theoretically, these ABPs could also be used to detect active CrMCA-II *in vivo*. However, we were thus far unable to find suitable conditions for microscopy to differentiate the Cy5 signal between UVM4 and *crmca-ii* mutant, presumably due to low abundance of the mature protease in the cells. An alternative explanation is poor permeability of the Cy5 probe to the pool(s) of active CrMCA-II. Therefore, further optimization of the chemical structure of these ABPs to improve permeability, as well as screening for physiologically relevant conditions facilitating CrMCA-II activation are both required for live imaging of CrMCA-II activity in a cell.

### PM localization and the thermoprotective role of CrMCA-II

MCAs feature diverse subcellular localization patterns, but most are nucleocytoplasmic and can be found in both soluble (cytosolic) and insoluble (protein aggregates and endoplasmic reticulum) protein fractions (Huh, 2022). Localization of CrMCA-II on PM is thus uncommon and therefore intriguing. The only other known MCA with similar localization validated experimentally is a *Trypanosoma brucei* TbMCA-Id/MCA4 of type I — a pseudopeptidase lacking catalytic Cys (Proto et al., 2011). Both CrMCA-II and TbMCA-Id/MCA4 are devoid of transmembrane domain(s), pointing to their strong affinity to lipids and/or other membranal proteins. Accordingly, TbMCA-Id/MCA4 has been shown to undergo *N*-myristoylation and palmitoylation *in vivo* to become associated with the internal surface of flagellar membrane (Proto et al., 2011). Being devoid of a *N*-myristoylation motif, the Cr-MCA-II sequence yet contains a number of predicted palmitoylation motifs (Ning et al., 2020; data not shown), which might contribute to the PM localization of the protein.

Interestingly, the presence of transmembrane domains is a widespread feature among orthocaspases and type I MCAs in prokaryotes (Klemenčič and Funk, 2018; La et al., 2022). Furthermore, transmembrane domains are predicted to occur (DeepTMHMM, Hallgren et al., 2022), albeit with much lower frequency, in type I MCAs from unicellular green algae, including *Auxenochlorella protothecoides* 0710 and *Picochlorum sp*. It thus seems likely that direct association with the PM is an ancient feature of C14 proteases, which was preserved by prokaryotes and some single celled eukaryotes but lost during the evolution of multicellularity. The latter assumption, however, is not completely true, since animal caspase-8 was shown to dimerize and become activated upon assembly of death inducing signaling complex (DISC) at the PM during initiation of the extrinsic pathway of apoptosis (Feig et al., 2007). Apart from the proteolytic role in apoptosis, caspase-8 promotes adhesion and Erk signaling in neuroblastoma cells *via* recruiting PM-associated Src tyrosine kinase (Finlay and Vuori, 2007). Notably, this function of caspase-8 is independent of its catalytic activity, resembling the non-catalytic role of CrMCA-II revealed in the current study. Nonetheless, in contrast to CrMCA-II, both procaspase-8 (zymogen) and mature caspase-8 enzyme are present only in the cytosolic S-100 fraction and are absent in the light and heavy membrane fractions (Breckenridge et al., 2002; Figure 5B).

As only the full-length zymogen but not processed CrMCA-II is present at the PM, while the cytosol and the endomembranes contain both types of proteoforms (Figure 5B), there must be a tight spatial control over protease activity of CrMCA-II in a cell. Thus, PM could serve as a compartment sequestering zymogen, whereas CrMCA-II protease maturation and substrate cleavage might take place in the cytoplasm. Given the critical requirement of Ca^2+^ for CrMCA-II activation and that the endoplasmic reticulum (ER) is a Ca^2+^ storage seat with a high concentration (Stael et al., 2011), it is reasonable to assume that the proteolytic function of CrMCA-II (if any) is fulfilled in or around the ER when Ca^2+^ is released. However, testing a panel of knockout and complementation strains under varied environmental settings failed to identify conditions wherein Chlamydomonas growth was dependent on the protease activity of CrMCA-II.

The PM plays a fundamental role in intra- and intercellular communication and in response to adverse environmental conditions, serving also as a crucial component of cell repair machinery (Horn and Jaiswal, 2019; Jaillais and Ott, 2019). The non-proteolytic role of the PM-localized CrMCA-II zymogen in HS could be related to its ability to modulate PM fluidity, alleviating hypofluidity under normal conditions and, correspondingly, preventing a large-amplitude hyperfluidity response to high temperatures (see model in Figure 6). Indeed, changes in PM fluidity have been shown to play a causative role in or correlate with various types of cell death in animals (summarized in Tekpli et al., 2013). In Chlamydomonas, treatment with membrane rigidifier DMSO decreases cell viability (Haire et al., 2018). Accordingly, the hypofluid PM in *crmca-ii* strains may fail to support transient lipid mobility after heat injury, undermining cell repair and leading to increased cell death (Horn and Jaiswal, 2019).

**Figure 6.**
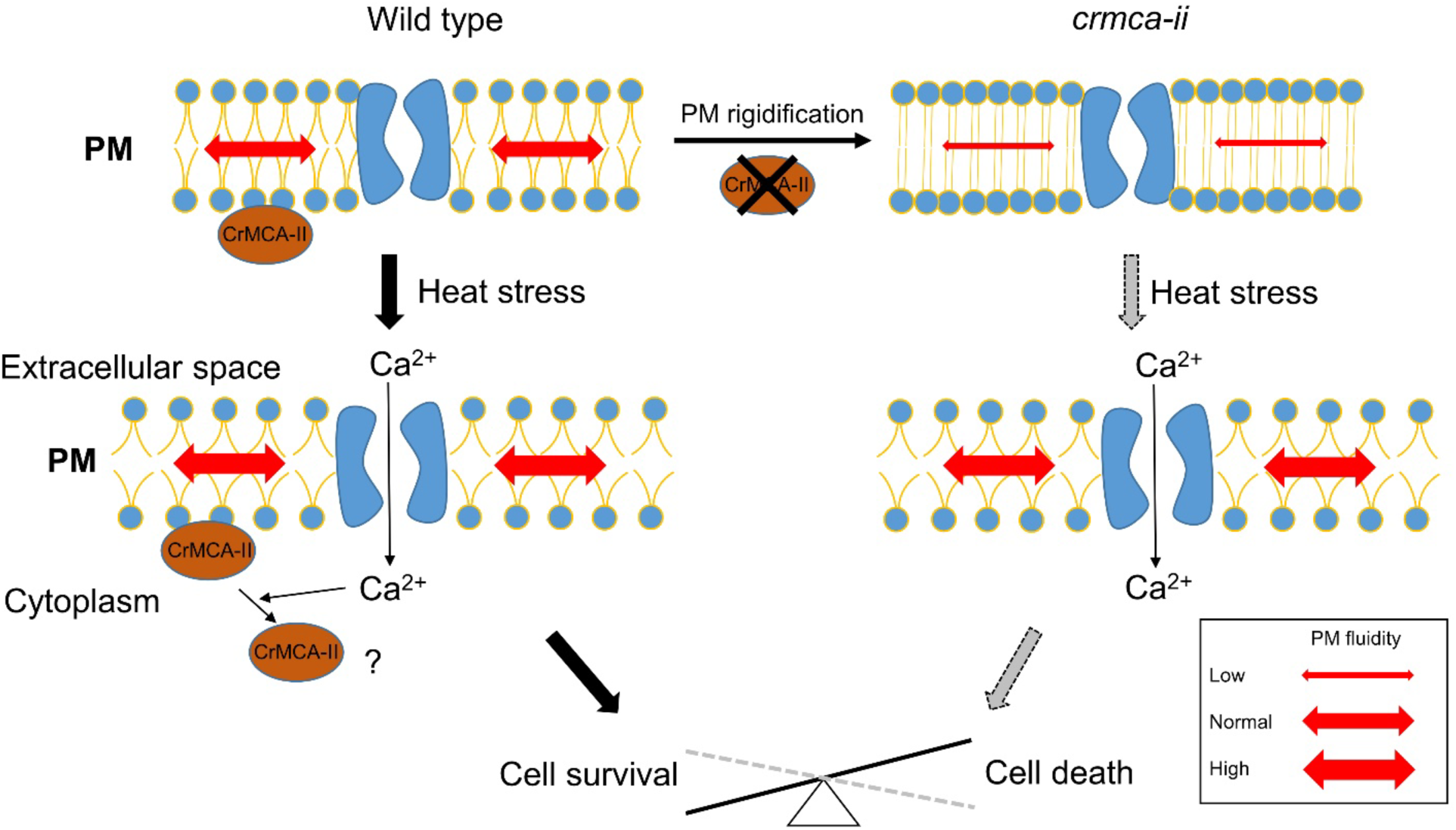
Hypothetical model for the CrMCA-II-dependent regulation of thermotolerance. Under favorable growth conditions, CrMCA-II is preferentially present as a zymogen attached to the PM and preventing its rigidification through an as yet unknown mechanism. HS elevates the PM fluidity, an increase becoming more dramatic and disrupting viability in the *crmca-ii* cells with rigidified PM, accounting for their decreased thermotolerance. An increase in the PM fluidity under HS stimulates influx of Ca^2+^ (Schroda et al., 2015) which, in turn, mediates the PM-to-cytoplasm translocation of CrMCA-II. The role of the cytoplasmic CrMCA-II under HS and, in particular, whether it contributes to cytoprotection remains unknown.

In contrast to animals (Shuba, 2021), heat response in plants remains poorly characterized. It is postulated that both streptophytes and chlorophytes sense heat *via* an increase in PM fluidity causing activation of the cyclic nucleotide-gated ion channels and controlled entry of extracellular Ca^2+^ to the cytosol, which further leads to the heat shock factor (HSF)-mediated transcriptional response and biosynthesis of necessary proteins, such as heat shock proteins (HSPs) (Mittler et al., 2012; Cano-Ramirez et al., 2021; Guihur et al., 2022). We initially thought that the compromised thermotolerance of *crmca-ii* strains was associated with aberrant HS sensing at the level of HSF and HSP production, but did not observe any differences in the transcript levels of HSFs and major HSPs known to execute HS response in Chlamydomonas (Zhang et al., 2022) between UVM4 and *crmca-ii* (data not shown). We therefore conclude that the mechanism of CrMCA-II-dependent thermoprotection is restricted to the regulation of physico-chemical properties of the PM *per se* affecting its fluidity, bilayer stability and hence functionality upon temperature shift (Horváth et al., 2008).

There are multiple factors underlying changes in PM fluidity, such as the alterations in the conformation of integral membrane proteins, density and size of lipid rafts, and saturation level of phospholipids (Desai and Miller, 2018; Pozza et al., 2022). Which of these factors are controlled or mediated by CrMCA-II will be addressed in our future work that will include characterization of CrMCA-II interactome and, specifically, zymogen-containing megadalton-scale complex (Figure 1G), as well as lipidomics analysis of the *crmca-ii* strains. Intriguingly, CrMCA-II is predicted to harbour several intrinsically disordered segments, spanning the non-catalytic regions of the protein (data not shown). Moreover, CrMCA-II has been recently identified as a part of a heat-resistant intrinsically disordered proteome of Chlamydomonas (Zhang et al., 2018). This observation together with our findings that the heat treatment stabilizes the zymogen and inhibits proteolytic activity of rCrMCA-II (Figure 4C-E) reinforce genetic evidence for a non-proteolytic role of rCrMCA-II in thermotolerance.

Interestingly, HS induces PM-to-cytoplasm translocation of CrMCA-II, in a Ca^2+^ influx-dependent manner (Figures 5C-E and 6). According to recent structural data obtained for Arabidopsis AtMCA-IIa/MC4 (Zhu et al., 2020), Ca^2+^ could bind to the high affinity-binding pocket of CrMCA-II (Asp95, Asp97, Glu98, and Asp100; Supplemental Figure S3), and impose structural changes causing its detachment from the PM. Whether PM-to-cytoplasm translocation of CrMCA-II has any functional role in HS response remains an open question.

In conclusion, CrMCA-II, - a single type II MCA of Chlamydomonas, - confers thermotolerance *via* a mechanism governing PM fluidity, which is completely independent of its protease activity. In this respect, the non-catalytic mode of CrMCA-II action is reminiscent of that of ScMCA-I/MCA1/YCA1, a single MCA encoded by *Saccharomyces cerevisiae* genome, in suppressing protein aggregate formation (also under HS) and replicative aging, albeit catalytic Cys276 mutant of the yeast MCA1 could only partially rescue *mca1* deletion phenotypes (Lee et al., 2010; Hill et al., 2014). Taken together, the findings in budding yeast and Chlamydomonas suggest an evolutionary scenario wherein the ancient primordial role of the MCAs was to carry out quality control both in the cytoplasm (e.g., to counteract protein aggregation) and at the cell surface (e.g., to maintain membrane fluidity) for fitness and survival of unicellular organisms in fluctuating environments. Complete or partial independence of intricately-regulated proteolytic activity could provide additional robustness to the proposed role of MCAs.

## Methods

### Strains and culture conditions

Unless stated otherwise, UV mutagenesis strain 4 (UVM4), with high efficiency to overexpress foreign genes (Neupert et al., 2009), was used as a control strain. The Chlamydomonas cells were grown in tris-acetate-phosphate (TAP) medium at 23°C and a light-dark cycle (LD16:8). The cells were harvested at a density of 1-2×10^7^ cells mL^-1^.

### Phylogenetic analysis

The MCA sequences of green algae and higher plants were obtained from Phytozome 13 (https://phytozome.jgi.doe.gov/pz/portal.html). The MEGA X (Kumar et al., 2018) was used to perform alignment with embedded Muscle program and phylogenetic analysis with default setting for neighbour-joining method.

### rCrMCA-II expression and purification

rCrMCA-II was expressed using a codon-optimized construct in *E. coli* and then purified through affinity chromatography and size exclusion chromatography (SEC) as described (Sabljić et al., 2022).

### Protease activity assay of rCrMCA-II

The proteolytic activity of 5 ng purified rCrMCA-II was measured in the reaction buffer (*viz.* optimal buffer conditions: 50 mM Tris pH 7.5, 25 mM NaCl, 20 mM CaCl_2_, 0.1% CHAPS, and 7.5 mM DTT), with 50 µM VRPR-AMC as a substrate, unless stated otherwise. Fluorescence intensity was measured every minute using BMG Labtech POLARstar Omega Microplate Readers at excitation/emission 360–380 nm/440–460 nm. The activity was determined based on the slope of at least 10 recordings.

### rCrMCA-II protease activity assay in native electrophoresis

To perform native electrophoresis, a 4-20% gradient stain-free gel was loaded with protein aliquots that had been incubated with 7.5 mM DTT for 15 min at 4°C. Controls were treated the same way, but without DTT. The gel was then run at 150 V, for 45 min in Tris/Glycine buffer. After gel running, 3 mL of the DTT-free reaction buffer was added on top of the gel and fluorescence was detected using a BioRad Gel Doc EZ Imager programmed for ethidium bromide.

### SEC of Chlamydomonas total protein extract

Cells were harvested in the late log phase from 100 mL TAP medium and suspended in 1mL extraction buffer (50 mM Tris pH 7.5, 25-100 mM NaCl, 1 mM EDTA, 1% glycerol, 1 µM Aprotinin, 10 µM Pepstatin A, 1 mM PMSF and 2 mM Pefabloc SC). The cell mixture was sonicated for 3 min by 10-sec pulses, with 5-sec intervals and the cell debris was pelleted at 17,000 *g* for 20 min. The resulting crude cell extract was filtered through a 0.45-µm filter before loading onto a HiLoad 16/60 Superdex 200 pg column (GE Healthcare) at 4°C. A buffer containing 50 mM Tris (pH 7.5) and 25 mM NaCl was used for equilibration and elution, and elution profiles were recorded at 280 nm. Fractions of 750 µL were collected, precipitated with 90% methanol (Bychkov et al., 2011), and dissolved in 40 µL 1 × SDS sample buffer. Immunoblots were performed after separating 10 µL of each fraction by SDS-PAGE in a 4-20% gel. A set of protein standards was used as a reference for calibration and determination of the molecular mass of CrMCA-II protein species in the samples.

### Chemical reagents

All chemicals used for the synthesis of substrates and activity-based probes (ABPs) were purchased from commercial suppliers and used without purification unless otherwise noted. The Rink amide RA resin (loading 0.48 mmol g^-1^) was used for the synthesis of ACC-labeled substrates, and the 2-chlorotrityl chloride resin (1.59 mmol g^-1^, 100-200 mesh) was used for the synthesis of peptides that were further converted into ABPs (both resins were from Iris Biotech GmbH, Germany). Fmoc-protected amino acids (all > 98% pure) were purchased from various suppliers: Iris Biotech GmbH, Creosalus, P3 BioSystems, QM Bio, Bachem. N,N-diisopropylethylamine (DIPEA, peptide grade), diisopropylcarbodiimide (DICI, peptide grade), piperidine (PIP, peptide grade), and trifluoroacetic acid (TFA, purity 99%) were all from Iris Biotech, GmbH. 2,4,6-trimethylpyridine (2,4,6-collidine, peptide grade), triisopropylsilane (TIPS, purity 99%), 2,2,2-trifluoroethanol (TFE), anhydrous tetrahydrofuran (THF), hydrogen bromide (30% wt. in AcOH), 4-methylmorpholine (NMM), isobutylchloroformate (IBCF), and 2,6-dimethylbenzoic acid (2,6-DMBA) were all purchased from Sigma Aldrich. N-hydroxybenzotriazole (HOBt, monohydrate) was from Creosalus. HATU and HBTU (both peptide grade) were from ChemPep Inc.. N,N′-dimethylformamide (DMF, peptide grade) and acetonitrile (ACN, HPLC pure) were from WITKO Sp. z o.o., Poland. Methanol (MeOH, pure for analysis), dichloromethane (DCM, pure for analysis), diethyl ether (Et_2_O, pure for analysis), acetic acid (AcOH, 98% pure) and phosphorus pentoxide (P_2_O_5_, 98% pure) were from POCh, Poland. Cyanine-5 NHS was purchased from Lumiprobe. Diazomethane was generated according to the Aldrich Technical Bulletin (AL-180) protocol.

### Characterization of rCrMCA-II P4-P2 substrate specificity using HyCoSuL

To determine the CrMCA-II specificity in the P4-P2 positions, we used the P1-Arg HyCoSuL library (Poreba et al., 2017). P4, P3, and P2 sub-libraries were each screened at 100 µM final concentration in 100 µL final volume with rCrMCA-II. The active enzyme was in the 2-10 nM range, depending on the sub-library used. The screening time was 30 min, but for each substrate cleavage only the linear portion of the progression curve (10-15 min) was used for the analysis (RFU sec^-1^ calculation) to avoid problems of substrate consumption. Screening of each library was performed at least in triplicate, and the average value was used to create the rCrMCA-II specificity matrix (S.D. for each substrate was below 15%). The hydrolysis rate for the best recognized amino acid at each position was set to 100%, and other amino acids were adjusted accordingly.

### Synthesis and screening of individual optimized substrates

ACC-labeled tetrapeptide substrates were synthesized and purified as described elsewhere (Poręba et al., 2014). Substrate hydrolysis at the concentration of 100 µM was carried out for 30 min. rCrMCA-II concentration was 2.1 nM. Each experiment was performed in triplicate and the rate of substrate hydrolysis (RFU sec^-1^) was presented as an average value (S.D. for each substrate hydrolysis was below 15%).

### Enzyme kinetics

All kinetics experiments were performed using a Gemini XPS (Molecular Devices) plate reader operating in the fluorescence kinetic mode using 96-well plates (Corning®, Costar®). ACC-labeled fluorescent substrates were screened at 355 nm (excitation) and 460 nm (emission) wavelengths under optimal buffer conditions. Buffer was prepared at an ambient temperature and all kinetics measurements were performed at 37°C. The kinetic parameters (k_cat_, K_M_, k_cat_/K_M_) for ACC-labeled substrates were determined using Michaelis-Menten kinetics according to the protocol described by Poreba et al. (2014). All experiments were performed at least in triplicate and the results were presented as average values (S.D. for all tested substrates was below 15%).

### Synthesis of fluorescent ABPs

Cy5-labeled irreversible ABPs probes for rCrMCA-II were synthesized and purified as described previously (Poreba et al., 2018). In brief, Boc-Ahx-peptide-COOH was synthesized on solid support using 2-chlorotrityl chloride resin and used without further purification. In parallel, the Boc-Arg(Boc)_2_-AOMK warhead was synthesized through generation of diazomethane and reaction with mixed anhydrides (Boc-Arg(Boc)_2_-OH → Boc-Arg(Boc)_2_-CH_2_N_2_), following by the conversion of the crude product into bromomethyl ketone (Boc-Arg(Boc)_2_-BMK) and, finally, into acyloxymethyl ketone (Boc-Arg(Boc)_2_-AOMK). Next, the crude product was de-protected in TFA/DCM mixture, and the NH_2_-Arg-AOMK was coupled with Boc-(linker)-peptide-COOH to yield Boc-(linker)-peptide-Arg-AOMK. The product was purified by HPLC, and after Boc de-protection, it was labeled with Cy5 to obtain Cy5-Ahx-peptide-Arg-AOMK. The final products were purified by HPLC, analyzed via HR-MS and HPLC and dissolved in DMSO (10 mM).

### ABP binding assay

Each reaction contained 0.4 µM rCrMCA-II or rCrMCA-II^C141A^ and indicated concentration of the ABP co-incubated for 5 min. The reaction was stopped by adding SDS sample buffer. Two-hundred ng purified protein was loaded onto an SDS-PAGE gel, which was then scanned at 700 nm using an Odyssey infrared scanner (LI-COR Biosciences). Total protein was visualized after staining with Oriole fluorescent gel stain solution (Bio-Rad) for 1.5-2 h.

### Genome editing

A Crispr/Cas9 based targeted insertional mutagenesis approach (Picariello et al., 2020) was employed to knockout the *CrMCA-II* gene. A guiding RNA was designed for targeting the exon region flanking the sequences encoding the catalytic dyad to disrupt the *CrMCA-II*. Electroporation was used to introduce a donor DNA comprising of 40 bp flanking sequences, *Hsp70/RBCS2* promoter*, AphVIII* gene, *RBCS2* terminator, and the ribonucleoprotein complex. The resulting transformants were screened with colony PCR and confirmed by immunoblotting with *α*-CrMCA-II.

### Genomic DNA isolation

Ten mL of cells grown in TAP were harvested and resuspended in 150 µL H_2_O and 300 µL of SDS-EB buffer (2% SDS, 400 mM NaCl, 40 mM EDTA, 100 mM Tris-HCl, pH 8.0). The cell suspension was mixed with 350 µL of phenol:chloroform:isoamyl alcohol (25:24:1) by inverting for few min, which led to the separation of two phases after centrifugation at 2,000 *g* for 5 min. The upper aqueous phase was extracted using 300 µL chloroform:isoamyl alcohol (24:1). The resulting aqueous phase was mixed with two volumes of absolute ethanol and stored at −80°C for 30 min. The genomic DNA was pelleted by centrifugation at 17,000 *g* for 10 min and washed twice with 200 µL 70% ethanol. The genomic DNA was then air-dried and dissolved in H_2_O.

### RNA isolation and quantitative PCR (qPCR)

RNA was isolated using Spectrum™ Plant Total RNA Kit (Sigma-Aldrich) according to the manufacturer’s instructions. DNAse was added during isolation to remove any remaining genomic DNA. For qPCR analysis, 200 ng of genomic DNA or 12.5 ng complementary DNA of total RNA was used in each reaction. The relative abundances of target genes were calculated using the 2^-ΔΔCt^ method (Livak and Schmittgen, 2001) with RACK1 (Mus et al., 2007) as a reference for normalization. All used primers are listed in Supplemental Table S2.

### Proteolytic activity assay of Chlamydomonas cell lysates

Ten ml of cells at the density of ∼ 1× 10^7^ cells mL^-1^ were harvested and stored at −80°C. Before extraction, the cells were thawed on ice for 5 min. Each sample was then suspended in 150 µL of extraction buffer (50 mM Tris pH 7.5, 25-100 mM NaCl, 1 mM EDTA, 1% glycerol, 1 µM aprotinin, 10 µM pepstatin A, 1 mM PMSF and 2 mM Pefabloc SC) and sonicated for 30 sec by 10-sec pulses, with 5-sec intervals, and the cell debris was pelleted at 7,800 *g* for 10 min. The resulting supernatant (cell lysate) was used for the assay. The proteolytic activity was measured in 50 µL extraction buffer supplemented with 0.1% CHAPS, 7.5 mM DTT, 20 mM CaCl_2,_ 50 µM of either VRPR-7-amino-4-methylcoumarin (AMC) or HRTR-ACC, and 1 µL cell lysate. The fluorescent signal was detected by POLARstar Omega Plate Reader (BMG LABTECH).

### Yeast Two-Hybrid (Y2H) assay

The Y2H experiments were performed as previous described (Zou et al., 2020). The cDNA sequences of CrMCA-II and serpin were cloned into AD or/and BD vector. The BD-CrMCA-II vector was used as a template to construct BD-CrMCA-II-C141A.

### Chlamydomonas subcellular fractionation

We followed previously published procedures (Norling et al., 1996; Herbik et al., 2002) with some modifications (see Supplemental Figure S13). Briefly, cells were harvested at 4,000 *g* for 5 min and snap-frozen in liquid nitrogen. One to 2 g cells were mixed with 40 mL extraction buffer (250 mM sucrose, 50 mM HEPES pH 7.5, 1 mM EGTA, 1 mM KCl, 0.6% w/v PVPP, 0.1 mM PMSF and 2.5 mM DTT) for 5 min on ice before sonicating twice for 10 min on ice. The unbroken cells, chloroplasts and cell debris were removed by centrifugation for 3 min at 1,200 *g*. The first supernatant (S1) was centrifuged for 5 min at 13,000 *g* to remove the pellet containing mainly mitochondria and thylakoids, and to collect the second supernatant (S2) containing low density membranes (LDMs) of Golgi, ER, tonoplast, as well as PM. The S2 was centrifuged for 2 h at 68,000 *g* at 4°C to separate into the LDM-containing pellet (P3) and a supernatant (S3). The P3 was re-suspended in a maximum of 2.5 mL re-suspension buffer (250 mM sucrose, 5 mM potassium phosphate pH 7.8, and 5 mM DTT), and the PM vesicles were separated from the endomembranes by two-phase partitioning system. The P3 suspension (2.25 g) was loaded onto an ice-cold polymer mixture (6.75 g) comprised of 6.4% dextran T500, 6.4% polyethyleneglycol (PEG, MW 4,000), 250 mM sucrose, 5 mM K-phosphate (pH 7.8) and 5 mM KCl. After centrifugation at 2,000 *g* for 10 min, the mixture separated into two phases. The upper PEG phase, containing the PM, was further partitioned against a fresh lower phase to increase the purity of the PM fraction. Subsequently, the PEG and dextran phases were diluted five-fold with the dilution buffer (0.33 M sucrose, 5 mM MOPS pH 7.0, 1 mM EDTA and 2 mM DTT). The PM proteins and the endomembrane proteins were pelleted from the PEG phase and the dextran phase, respectively by centrifugation at 68,000 *g* for 1 h at 4°C. The endomembrane and the PM proteins were re-suspended in dilution buffer for immunoblotting.

### Protein extraction and immunoblotting

Two ml cells in late log phase (1-2×10^7^ cells mL^-1^) were harvested by centrifugation at 17, 000 *g* for 1 min and were snap-frozen in liquid nitrogen. The cell pellet was dissolved in 200 µL 1×SDS sample buffer (50 mM Tris-Cl pH 7.5, 100 mM DTT, 2% (w/v) SDS, 0.1% (w/v) Bromophenol Blue, 10% (w/v) glycerol) and incubated at 70°C for 5 min. The samples were vortexed at the highest speed and then centrifuged at 17,000 *g* for 10 min. The resulting supernatant was used for immunoblotting. The *α*-CrMCA-II was generated by Agrisera using peptide antigen characterized in Supplemental Figure S11A. The *α*-H^+^ ATPase was purchase from Agrisera. Primary antibody dilutions for *α*-CrMCA-II and *α*-H^+^ ATPase were 1:1,000 and 1:5,000, respectively. A secondary anti-rabbit antibody was used at a dilution of 1:10,000.

### Cell death measurements

Cells in log phase (∼5×10^6^ cells mL^-1^) were subjected to HS at 42°C for 1 h and then cooled down to room temperature before treated with Fluorescein Diacetate (FDA, final concentration 2 µg mL^-1^; Minina et al., 2013). The FDA-treated cells were loaded into hemocytometer and photographed. When excited at 480 nm, living cells appeared green, while dead cells appeared red. Cell death was calculated using the following formula: cell death (%) = dead cells / (dead cells + living cells) *100%.

### Live imaging of CrMCA-II

The *CrMCA-II* gene was cloned into the MoClo toolkit (Crozet et al., 2018). Subsequently, a plasmid containing *PsaD* promoter, the genomic sequence of *CrMCA-II*, the *mVenus* gene and the *PsaD* terminator was assembled to overexpress CrMCA-II-mVenus. The transformants at log phase were screened and imaged by confocal microscopy (Zeiss LSM 780), with the excitation and emission at 514 nm and 540-580 nm, respectively.

### Fluorescence recovery after photo-bleaching (FRAP)

Due to the high sensitivity to photobleaching at 42°C HS resulting in cell damage, the temperature of HS for the FRAP analysis was reduced to 39°C. The cells were labeled with DiOC6(3) (Invitrogen, final concentration: 0.05 µg/ml) for 1 min at room temperature, followed by 2 washes with fresh TAP medium. The labeled PM was observed by confocal microscopy (LSM 780, Carl Zeiss), with the excitation and emission at 488 nm and 501-570 nm, respectively. FRAP analysis was conducted as described (Gutierrez-Beltran et al., 2021), with the following modifications: 60 iterations, 1 sec per frame, and 75% transmittance with the 488-nm laser lines of the argon laser. Pre- and post-bleach scans were performed using the laser power at 0.5% to 2% transmittance for 488 nm and 0% for all other laser lines.

## Acknowledgements

This work was supported by the Knut and Alice Wallenberg Foundation (grants 2018.0026 to P.V.B. and J.S and 2021.0071 to S.S.) and (grant 2019-01565 to A.N.D.). We would like to thank Dr. Marina Klemenčič (University of Ljubljana) for sharing a plasmid overexpressing CrMCA-II and for fruitful discussions, Dr. George B. Witman and Dr. Yuqing Hou (both from University of Massachusetts Chan Medical School) for advice on genome editing, Dr. Ralph Bock (Max Planck Institute of Molecular Plant Physiology) for sharing algal strain UVM4, and Dr. Anna Maria Eisele (SLU, Uppsala) for fruitful discussions.

## Author contributions

Y.Z. and P.V.B. designed the study with input from I.S., A.N.D., S.S., M.P., and J.S. Y.Z., I.S., M.P. and P.V.B. wrote the manuscript. Y.Z., I.S., N.H., A.N.D., A.Å., and L.S.T. performed experiments. M.D. contributed new analytic tools.

## References

Becker, B., and Marin, B. (2009). Streptophyte algae and the origin of embryophytes. Ann Bot 103, 999–1004.

Bemer, K. (1985). Summary of green plant phylogeny and classification. Cladistics 1, 369–385.

Bollhöner, B., Zhang, B., Stael, S., Denancé, N., Overmyer, K., Goffner, D., Van Breusegem, F., and Tuominen, H. (2013). *Post mortem* function of AtMC9 in xylem vessel elements. New Phytol 200, 498–510.

Bozhkov, P.V., Suarez, M.F., Filonova, L.H., Daniel, G., Zamyatnin, A.A., Rodriguez-Nieto, S., Zhivotovsky, B., and Smertenko, A. (2005). Cysteine protease mcII-Pa executes programmed cell death during plant embryogenesis. Proc Natl Acad Sci USA 102, 14463–14468.

Breckenridge, D.G., Nguyen, M., Kuppig, S., Reth, M., and Shore, G.C. (2002). The procaspase-8 isoform, procaspase-8L, recruited to the BAP31 complex at the endoplasmic reticulum. Proc Natl Acad Sci USA 99, 4331–4336.

Bychkov, E.R., Ahmed, M.R., Gurevich, V.V., Benovic, J.L., and Gurevich, E.V. (2011). Reduced expression of G protein-coupled receptor kinases in schizophrenia but not in schizoaffective disorder. Neurobiol Dis 44, 248–258.

Cano-Ramirez, D.L., Carmona-Salazar, L., Morales-Cedillo, F., Ramírez-Salcedo, J., Cahoon, E.B., and Gavilanes-Ruíz, M. (2021). Plasma membrane fluidity: An environment thermal detector in plants. Cells 10, 2778.

Cerretti, D.P., Kozlosky, C.J., Mosley, B., Nelson, N., Ness, K.V., Greenstreet, T.A., March, C.J., Kronheim, S.R., Druck, T., Cannizzaro, L.A., Huebner, K., and Black, R.A. (1992). Molecular cloning of the interleukin-1β converting enzyme. Science 256, 97–100.

Choi, C.J., and Berges, J.A. (2013). New types of metacaspases in phytoplankton reveal diverse origins of cell death proteases. Cell Death Dis 4, e490.

Coll, N.S., Smidler, A., Puigvert, M., Popa, C., Valls, M., and Dangl, J.L. (2014). The plant metacaspase AtMC1 in pathogen-triggered programmed cell death and aging: functional linkage with autophagy. Cell Death Differ 21, 1399–1408.

Coll, N.S., Vercammen, D., Smidler, A., Clover, C., Van Breusegem, F., Dangl, J.L., and Epple, P. (2010). *Arabidopsis* type I metacaspases control cell death. Science 330, 1393–1397.

Conchou, L., Doumèche, B., Galisson, F., Violot, S., Dugelay, C., Diesis, E., Page, A., Bienvenu, A.-L., Picot, S., Aghajari, N., and Ballut, L. (2022). Structural and molecular determinants of Candida glabrata metacaspase maturation and activation by calcium. Commun Biol 5, 1158.

Crozet, P., Navarro, F.J., Willmund, F., Mehrshahi, P., Bakowski, K., and, et al. (2018). Birth of a photosynthetic chassis: A MoClo toolkit enabling synthetic biology in the microalga *Chlamydomonas reinhardtii*. Acs Synth Biol 7, 2074–2086.

Desai, A.J., and Miller, L.J. (2018). Changes in the plasma membrane in metabolic disease: impact of the membrane environment on G protein-coupled receptor structure and function. Br J Pharmacol 175, 4009–4025.

Feig, C., Tchikov, V., Schütze, S., and Peter, M.E. (2007). Palmitoylation of CD95 facilitates formation of SDS-stable receptor aggregates that initiate apoptosis signaling. EMBO J 26, 221–231.

Finlay, D., and Vuori, K. (2007). Novel noncatalytic role for caspase-8 in promoting Src-mediated adhesion and Erk signaling in neuroblastoma cells. Cancer Res 67, 11704–11711.

Graff van Creveld, S., Ben-Dor, S., Mizrachi, A., Alcolombri, U., Hopes, A., Mock, T., Rosenwasser, S., and Vardi, A. (2021). Biochemical characterization of a novel redox-regulated metacaspase in a marine diatom. Front Microbiol 12.

Guihur, A., Rebeaud, M.E., and Goloubinoff, P. (2022). How do plants feel the heat and survive? Trends Biochem Sci 47, 824–838.

Gutierrez-Beltran, E., Elander, P.H., Dalman, K., Dayhoff II, G.W., Moschou, P.N., Uversky, V.N., Crespo, J.L., and Bozhkov, P.V. (2021). Tudor staphylococcal nuclease is a docking platform for stress granule components and is essential for SnRK1 activation in Arabidopsis. EMBO J 40, e105043.

Gutman, B.L., and Niyogi, K.K. (2004). Chlamydomonas and Arabidopsis. A dynamic duo. Plant Physiol 135, 607–610.

Haire, T.C., Bell, C., Cutshaw, K., Swiger, B., Winkelmann, K., and Palmer, A.G. (2018). Robust microplate-based methods for culturing and in vivo phenotypic screening of *Chlamydomonas reinhardtii*. Front Plant Sci 9.

Hallgren, J., Tsirigos, K.D., Pedersen, M.D., Almagro Armenteros, J.J., Marcatili, P., Nielsen, H., Krogh, A., and Winther, O. (2022). DeepTMHMM predicts alpha and beta transmembrane proteins using deep neural networks. bioRxiv, 2022.2004.2008.487609.

Hander, T., Fernández-Fernández, Á.D., Kumpf, R.P., Willems, P., Schatowitz, H., Rombaut, D., Staes, A., Nolf, J., Pottie, R., Yao, P., Gonçalves, A., Pavie, B., Boller, T., Gevaert, K., Van Breusegem, F., Bartels, S., and Stael, S. (2019). Damage on plants activates Ca^2+^-dependent metacaspases for release of immunomodulatory peptides. Science 363, eaar7486.

Herbik, A., Haebel, S., and Buckhout, T.J. (2002). Is a ferroxidase involved in the high-affinity iron uptake in *Chlamydomonas reinhardtii*? Plant Soil 241, 1–10.

Hill, S.M., Hao, X.X., Liu, B.D., and Nystrom, T. (2014). Life-span extension by a metacaspase in the yeast Saccharomyces cerevisiae. Science 344, 1389–1392.

Hoebeeck, J., Speleman, F., and Vandesompele, J. (2007). Real-time quantitative PCR as an alternative to southern blot or fluorescence in situ hybridization for detection of gene copy number changes. In Methods Mol Biol (Protocols for Nucleic Acid Analysis by Nonradioactive Probes), E. Hilario and J. Mackay, eds (Totowa, NJ: Humana Press), pp. 205–226.

Horn, A., and Jaiswal, J.K. (2019). Structural and signaling role of lipids in plasma membrane repair. In Curr Top Membr, L.O. Andrade, ed (Academic Press), pp. 67–98.

Horváth, I., Multhoff, G., Sonnleitner, A., and Vígh, L. (2008). Membrane-associated stress proteins: More than simply chaperones. Biochim Biophys Acta Biomembr 1778, 1653–1664.

Huh, S.U. (2022). Evolutionary diversity and function of metacaspases in plants: Similar to but not caspases. Int J Mol Sci 23, 4588.

Jaillais, Y., and Ott, T. (2019). The nanoscale organization of the plasma membrane and its importance in signaling: A proteolipid perspective. Plant Physiol 182, 1682–1696.

Julien, O., and Wells, J.A. (2017). Caspases and their substrates. Cell Death Differ 24, 1380–1389.

Kato, D., Boatright, K.M., Berger, A.B., Nazif, T., Blum, G., Ryan, C., Chehade, K.A.H., Salvesen, G.S., and Bogyo, M. (2005). Activity-based probes that target diverse cysteine protease families. Nat Chem Biol 1, 33–38.

Kesavardhana, S., Malireddi, R.K.S., and Kanneganti, T.-D. (2020). Caspases in cell death, inflammation, and pyroptosis. Annu Rev Immunol 38, 567–595.

Klemenčič, M., and Funk, C. (2018). Structural and functional diversity of caspase homologues in non-metazoan organisms. Protoplasma 255, 387–397.

Klemenčič, M., and Funk, C. (2019). Evolution and structural diversity of metacaspases. J Exp Bot 70, 2039–2047.

Kumar, S., Stecher, G., Li, M., Knyaz, C., and Tamura, K. (2018). MEGA X: Molecular evolutionary genetics analysis across computing platforms. Mol Biol Evol 35, 1547–1549.

La, S.R., Ndhlovu, A., and Durand, P.M. (2022). The ancient origins of death domains support the ‘original sin’ hypothesis for the evolution of programmed cell death. J Mol Evol 90, 95–113.

Lam, E., and Zhang, Y. (2012). Regulating the reapers: activating metacaspases for programmed cell death. Trends in Plant Sci 17, 487–494.

Lee, R.E.C., Brunette, S., Puente, L.G., and Megeney, L.A. (2010). Metacaspase Yca1 is required for clearance of insoluble protein aggregates. Proc Natl Acad Sci USA 107, 13348–13353.

Li, Y., Xue, J., Wang, F.-Z., Huang, X., Gong, B.-Q., Tao, Y., Shen, W., Tao, K., Yao, N., Xiao, S., Zhou, J.-M., and Li, J.-F. (2022). Plasma membrane-nucleo-cytoplasmic coordination of a receptor-like cytoplasmic kinase promotes EDS1-dependent plant immunity. Nat Plants 8, 802–816.

Livak, K.J., and Schmittgen, T.D. (2001). Analysis of relative gene expression data using real-time quantitative PCR and the 2^−ΔΔCT^ method. Methods 25, 402–408.

Long, H., Zhang, F., Xu, N., Liu, G., Diener, D.R., Rosenbaum, J.L., and Huang, K. (2016). Comparative analysis of ciliary membranes and ectosomes. Curr Biol 26, 3327–3335.

MacKenzie, S.H., and Clark, A.C. (2012). Death by caspase dimerization. In Adv Exp Med Biol (Protein Dimerization and Oligomerization in Biology), J.M. Matthews, ed (New York, NY: Springer New York), pp. 55–73.

McConnell, E.W., Werth, E.G., and Hicks, L.M. (2018). The phosphorylated redox proteome of *Chlamydomonas reinhardtii*: Revealing novel means for regulation of protein structure and function. Redox Biol 17, 35–46.

McLuskey, K., and Mottram, Jeremy C. (2015). Comparative structural analysis of the caspase family with other clan CD cysteine peptidases. Biochem 466, 219–232.

McLuskey, K., Rudolf, J., Proto, W.R., Isaacs, N.W., Coombs, G.H., Moss, C.X., and Mottram, J.C. (2012). Crystal structure of a *Trypanosoma brucei* metacaspase. Proc Natl Acad Sci USA 109, 7469–7474.

Merchant, S.S., Prochnik, S.E., Vallon, O., Harris, E.H., Karpowicz, S.J., and et, a.l. (2007). The *Chlamydomonas* genome reveals the evolution of key animal and plant functions. Science 318, 245–250.

Minina, E.A., Coll, N.S., Tuominen, H., and Bozhkov, P.V. (2017). Metacaspases versus caspases in development and cell fate regulation. Cell Death Differ 24, 1314–1325.

Minina, E.A., Filonova, L.H., Sanchez-Vera, V., Suarez, M.F., Daniel, G., and Bozhkov, P.V. (2013). Detection and measurement of necrosis in plants. In Methods Protoc (Necrosis), K. McCall and C. Klein, eds (Totowa, NJ: Humana Press), pp. 229–248.

Minina, E.A., Staal, J., Alvarez, V.E., Berges, J.A., Berman-Frank, I., and et, a.l. (2020). Classification and nomenclature of metacaspases and paracaspases: No more confusion with caspases. Mol Cell 77, 927-929.

Mittler, R., Finka, A., and Goloubinoff, P. (2012). How do plants feel the heat? Trends Biochem Sci 37, 118–125.

Morris, J.L., Puttick, M.N., Clark, J.W., Edwards, D., Kenrick, P., Pressel, S., Wellman, C.H., Yang, Z., Schneider, H., and Donoghue, P.C.J. (2018). The timescale of early land plant evolution. Proc Natl Acad Sci USA 115, E2274–E2283.

Mus, F., Dubini, A., Seibert, M., Posewitz, M.C., and Grossman, A.R. (2007). Anaerobic acclimation in *Chlamydomonas reinhardtii*: anoxic gene expression, hydrogenase induction, and metabolic pathways. J Biol Chem 282, 25475–25486.

Neupert, J., Karcher, D., and Bock, R. (2009). Generation of *Chlamydomonas* strains that efficiently express nuclear transgenes. Plant J 57, 1140–1150.

Ning, W., Jiang, P., Guo, Y., Wang, C., Tan, X., Zhang, W., Peng, D., and Xue, Y. (2020). GPS-Palm: a deep learning-based graphic presentation system for the prediction of S-palmitoylation sites in proteins. Brief Bioinformatics 22, 1836–1847.

Niu, Y., and Xiang, Y. (2018). An overview of biomembrane functions in plant responses to high-temperature stress. Front Plant Sci 9.

Nordzieke, D.E., and Medraño-Fernandez, I. (2018). The plasma membrane: A platform for intra- and intercellular redox signaling. Antioxidants 7, 168.

Norling, B., Nurani, G., and Franzén, L.-G. (1996). Characterisation of the H^+^-ATPase in plasma membranes isolated from the green alga *Chlamydomonas reinhardtii*. Physiol Plant 97, 445–453.

Pérez-Pérez, M.E., Mauriès, A., Maes, A., Tourasse, N.J., Hamon, M., Lemaire, S.D., and Marchand, C.H. (2017). The deep thioredoxome in *Chlamydomonas reinhardtii*: New insights into redox regulation. Mol Plant 10, 1107–1125.

Picariello, T., Hou, Y., Kubo, T., McNeill, N.A., Yanagisawa, H.-a., Oda, T., and Witman, G.B. (2020). TIM, a targeted insertional mutagenesis method utilizing CRISPR/Cas9 in *Chlamydomonas reinhardtii*. PloS One 15, e0232594.

Poreba, M., Salvesen, G.S., and Drag, M. (2017). Synthesis of a HyCoSuL peptide substrate library to dissect protease substrate specificity. Nat Protoc 12, 2189–2214.

Poreba, M., Rut, W., Vizovisek, M., Groborz, K., Kasperkiewicz, P., Finlay, D., Vuori, K., Turk, D., Turk, B., Salvesen, G.S., and Drag, M. (2018). Selective imaging of cathepsin L in breast cancer by fluorescent activity-based probes. Chem 9, 2113–2129.

Poręba, M., Szalek, A., Kasperkiewicz, P., and Drąg, M. (2014). Positional scanning substrate combinatorial library (PS-SCL) approach to define caspase substrate specificity In Methods Protoc (Caspases,Paracaspases, and Metacaspases), P. V. Bozhkov and G. Salvesen, eds (New York, NY: Springer New York), pp. 41–59.

Pozza, A., Giraud, F., Cece, Q., Casiraghi, M., Point, E., Damian, M., Le Bon, C., Moncoq, K., Banères, J.-L., Lescop, E., and Catoire, L.J. (2022). Exploration of the dynamic interplay between lipids and membrane proteins by hydrostatic pressure. Nat Commun 13, 1780.

Proto, W.R., Castanys-Munoz, E., Black, A., Tetley, L., Moss, C.X., Juliano, L., Coombs, G.H., and Mottram, J.C. (2011). Trypanosoma brucei metacaspase 4 is a pseudopeptidase and a virulence factor. J Biol Chem 286, 39914–39925.

Ross, C., Chan, A.H., Pein, J.B.v., Maddugoda, M.P., Boucher, D., and Schroder, K. (2022). Inflammatory caspases: Toward a unified model for caspase activation by inflammasomes. Annu Rev Immunol 40, 249–269.

Sabljić, I., Zou, Y., Klemenčič, M., Funk, C., Ståhlberg, J., and Bozhkov, P. (2022). Expression andpurification of the type II metacaspase from a unicellular green alga *Chlamydomonas reinhardtii*. In Methods Protoc (Plant Proteases and Plant Cell Death), M. Klemenčič, S. Stael, and P.F. Huesgen, eds (New York, NY: Springer US), pp. 13–20.

Saidi, Y., Finka, A., Muriset, M., Bromberg, Z., Weiss, Y.G., Maathuis, F.J.M., and Goloubinoff, P. (2009). The heat shock response in moss plants is regulated by specific calcium-permeable channels in the plasma membrane Plant Cell 21, 2829–2843.

Salguero-Linares, J., and Coll, N.S. (2019). Plant proteases in the control of the hypersensitive response. J Exp Bot 70, 2087–2095.

Schneider-Poetsch, T., Ju, J., Eyler, D.E., Dang, Y., Bhat, S., Merrick, W.C., Green, R., Shen, B., and Liu, J.O. (2010). Inhibition of eukaryotic translation elongation by cycloheximide and lactimidomycin. Nat Chem Biol 6, 209–217.

Schroda, M., Hemme, D., and Mühlhaus, T. (2015). The *Chlamydomonas* heat stress response. Plant J. 82, 466–480.

Shen, W., Liu, J., and Li, J.-F. (2019). Type-II metacaspases mediate the processing of plant elicitor peptides in *Arabidopsis*. Mol Plant 12, 1524–1533.

Shuba, Y.M. (2021). Beyond neuronal heat sensing: Diversity of trpv1 heat-capsaicin receptor-channel functions. Front Cell Neurosci 14.

Stael, S., Wurzinger, B., Mair, A., Mehlmer, N., Vothknecht, U.C., and Teige, M. (2011). Plant organellar calcium signalling: an emerging field. J Exp Bot 63, 1525-1542.

Stael, S., Sabljić, I., Audenaert, D., Andersson, T., Tsiatsiani, L., Kumpf, R.P., Vidal-Albalat, A., Lindgren, C., Vercammen, D., Jacques, S., Nguyen, L., Njo, M., Fernández-Fernández, Á.D., Beunens, T., Timmerman, E., Gevaert, K., Ståhlberg, J., Bozhkov, P.V., Linusson, A., Beeckman, T., and Van Breusegem, F. (2023). Structure-function analysis of a Ca^2+^-independent metacaspase reveals a novel proteolytic pathway for lateral root emergence. Proc Natl Acad Sci USA 120, e2303480120.

Stennicke, H.R., and Salvesan, G.S. (1997). Biochemical characteristics of caspases-3, −6, −7, and −8. J Biol Chem 272, 25719–25723.

Sundström, J.F., Vaculova, A., Smertenko, A.P., Savenkov, E.I., Golovko, A., Minina, E., Tiwari, B.S., Rodriguez-Nieto, S., Zamyatnin Jr, A.A., Välineva, T., Saarikettu, J., Frilander, M.J., Suarez, M.F., Zavialov, A., Ståhl, U., Hussey, P.J., Silvennoinen, O., Sundberg, E., Zhivotovsky, B., and Bozhkov, P.V. (2009). Tudor staphylococcal nuclease is an evolutionarily conserved component of the programmed cell death degradome. Nat Cell Biol 11, 1347–1354.

Tekpli, X., Holme, J.A., Sergent, O., and Lagadic-Gossmann, D. (2013). Role for membrane remodeling in cell death: Implication for health and disease. Toxicology 304, 141–157.

Thornberry, N.A., Bull, H.G., Calaycay, J.R., Chapman, K.T., Howard, A.D., Kostura, M.J., Miller, D.K., Molineaux, S.M., Weidner, J.R., Aunins, J., Elliston, K.O., Ayala, J.M., Casano, F.J., Chin, J., Ding, G.J.F., Egger, L.A., Gaffney, E.P., Limjuco, G., Palyha, O.C., Raju, S.M., Rolando, A.M., Salley, J.P., Yamin, T.-T., Lee, T.D., Shively, J.E., MacCross, M., Mumford, R.A., Schmidt, J.A., and Tocci, M.J. (1992). A novel heterodimeric cysteine protease is required for interleukin-1β processing in monocytes. Nature 356, 768–774.

Tsiatsiani, L., Van Breusegem, F., Gallois, P., Zavialov, A., Lam, E., and Bozhkov, P.V. (2011). Metacaspases. Cell Death Differ 18, 1279-1288.

Tsiatsiani, L., Timmerman, E., De Bock, P.-J., Vercammen, D., Stael, S., van de Cotte, B., Staes, A., Goethals, M., Beunens, T., Van Damme, P., Gevaert, K., and Van Breusegem, F. (2013). The *Arabidopsis* METACASPASE9 degradome. The Plant Cell 25, 2831–2847.

Uren, A.G., O’Rourke, K., Aravind, L., Pisabarro, M.T., Seshagiri, S., Koonin, E.V., and Dixit, V.M. (2000). Identification of paracaspases and metacaspases: Two ancient families of caspase-like proteins, one of which plays a key role in MALT lymphoma. Mol Cell 6, 961–967.

Van Opdenbosch, N., and Lamkanfi, M. (2019). Caspases in cell death, inflammation, and disease. Immunity 50, 1352–1364.

Vercammen, D., van de Cotte, B., De Jaeger, G., Eeckhout, D., Casteels, P., Vandepoele, K., Vandenberghe, I., Van Beeumen, J., Inze, D., and Van Breusegem, F. (2004). Type II metacaspases Atmc4 and Atmc9 of *Arabidopsis thaliana* cleave substrates after arginine and lysine. J Biol Chem 279, 45329–45336.

Vercammen, D., Belenghi, B., van de Cotte, B., Beunens, T., Gavigan, J.-A., De Rycke, R., Brackenier, A., Inzé, D., Harris, J.L., and Van Breusegem, F. (2006). Serpin1 of *Arabidopsis thaliana* is a suicide inhibitor for metacaspase 9. J Mol Biol 364, 625–636.

Wang, S., Xue, M., He, C., Shen, D., Jiang, C., Zhao, H., and Niu, D. (2021). AtMC1 associates with LSM4 to regulate plant immunity through modulating pre-mRNA splicing. Mol Plant Microbe Interact 34, 1423–1432.

Watanabe, N., and Lam, E. (2011). Calcium-dependent activation and autolysis of *Arabidopsis* metacaspase 2d. J Biol Chem 286, 10027–10040.

Wong, A.H.-H., Yan, C., and Shi, Y. (2012). Crystal structure of the yeast metacaspase Yca1. J Biol Chem 87, 29251–29259.

Yuan, J., Shaham, S., Ledoux, S., Ellis, H.M., and Horvitz, H.R. (1993). The C. elegans cell death gene *ced-3* encodes a protein similar to mammalian interleukin-1β-converting enzyme. Cell 75, 641–652.

Zhang, N., Mattoon, E.M., McHargue, W., Venn, B., Zimmer, D., Pecani, K., Jeong, J., Anderson, C.M., Chen, C., Berry, J.C., Xia, M., Tzeng, S.-C., Becker, E., Pazouki, L., Evans, B., Cross, F., Cheng, J., Czymmek, K.J., Schroda, M., Mühlhaus, T., and Zhang, R. (2022). Systems-wide analysis revealed shared and unique responses to moderate and acute high temperatures in the green alga *Chlamydomonas reinhardtii*. Commun Biol 5, 460.

Zhang, Y., Launay, H., Schramm, A., Lebrun, R., and Gontero, B. (2018). Exploring intrinsically disordered proteins in *Chlamydomonas reinhardtii*. Sci Rep 8, 6805.

Zhu, P., Yu, X.H., Wang, C., Zhang, Q., Liu, W., McSweeney, S., Shanklin, J., Lam, E., and Liu, Q. (2020). Structural basis for Ca^2+^-dependent activation of a plant metacaspase. Nat Commun 11, 1–38.

Zou, Y., Li, R., and Baldwin, I.T. (2020). ZEITLUPE is required for shade avoidance in the wild tobacco *Nicotiana attenuata*. J Integr Plant Biol 62, 1341–1351.

