## supplemental info for "Thermoprotection by a cell membrane-localized metacaspase in a green alga"

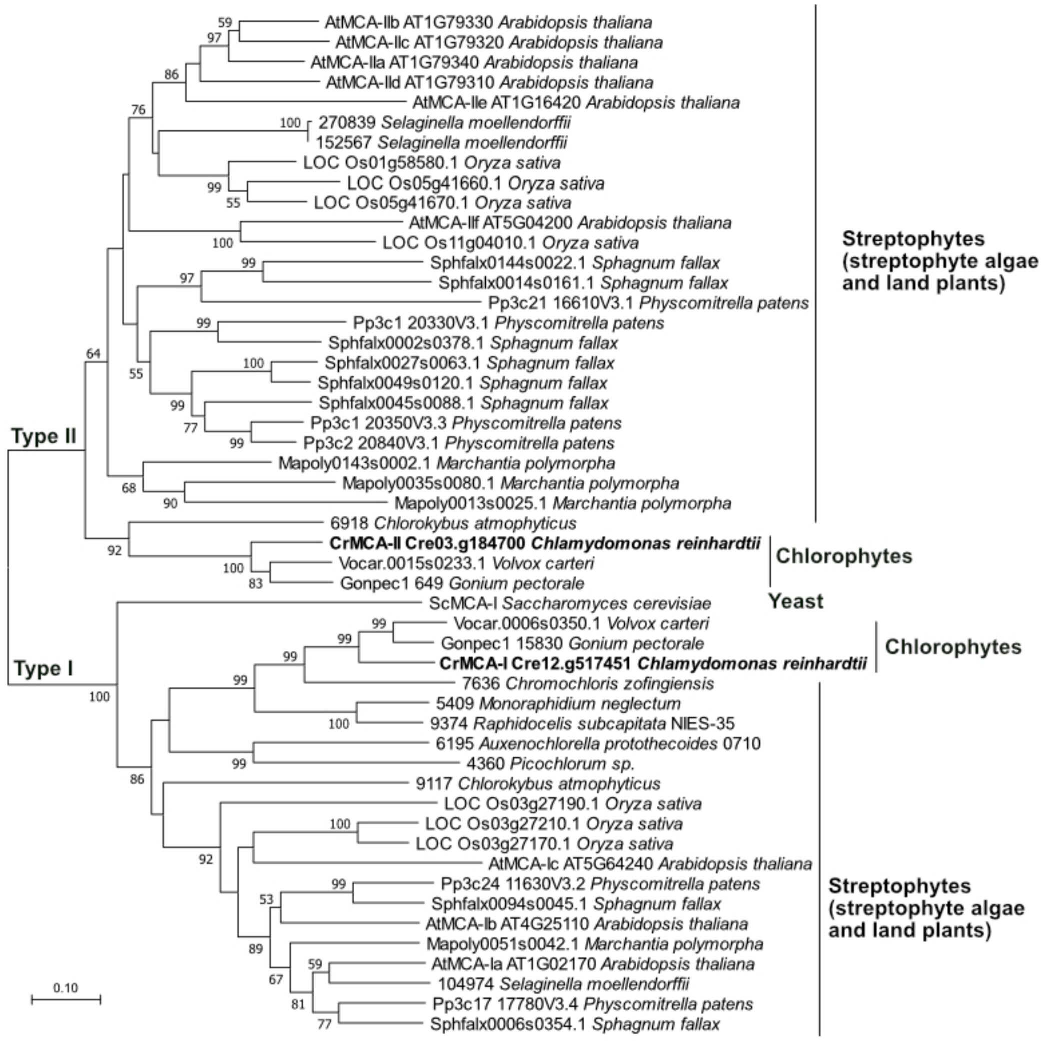


**Supplemental Figure S1** The phylogeny of MCAs from green plants (Chlorophytes and Streptophytes). Each sequence contains the accession number from the Phytozome 13 (https://phytozome-next.jgi.doe.gov/). The sequences from *Chlamydomonas reinhardtii* are highlighted in bold. A type I MCA from budding yeast is included. MEGA X was used for alignment and phylogenetic analysis, with default settings of the neighbor-joining method. A thousand replicates were run and the bootstrap values over 50 are displayed in percentage next to the branches. The scale bar, amino acid substitutions per site.

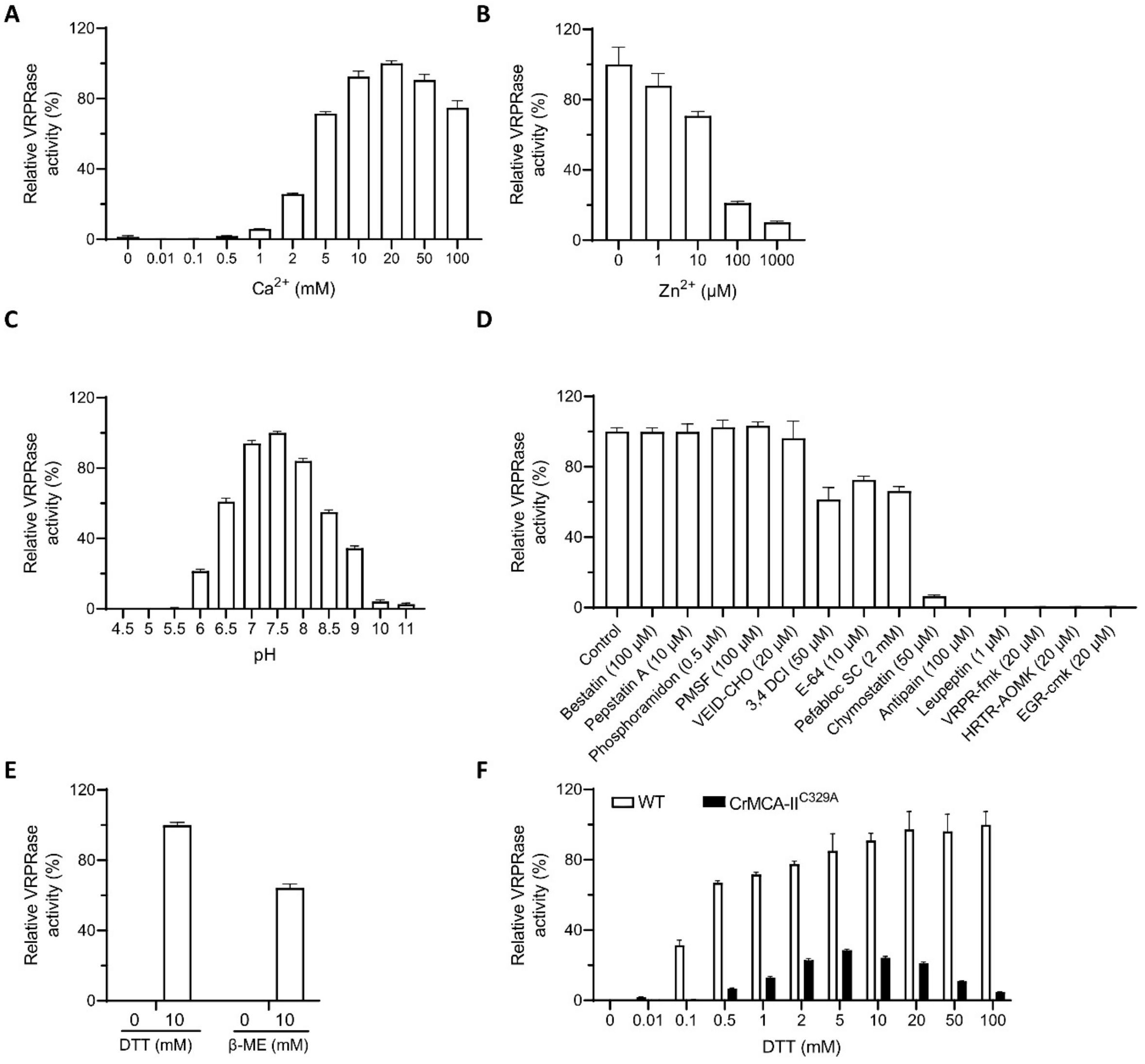


**Supplemental Figure S2** The biochemical characteristics of rCrMCA-II. Effects of $Ca^{2+}$ ($CaCl_2$) (**A**), zinc ($ZnCl_2$) (**B**), pH (**C**), protease inhibitors (**D**), and reducing agents (**E**) on proteolytic activity of rCrMCA-II under optimal buffer conditions (50 mM Tris pH 7.5, 25 mM NaCl, 20 mM $CaCl_2$, 0.1% CHAPS and 7.5 mM DTT, unless stated otherwise) using Ac-VRPR-AMC as a substrate. **F.** Activity of rCrMCA-II$^{C329A}$, as compared with a wild type (WT) rCrMCA-II (data from Fig. 1B), at increased concentration of DTT. Data on charts represent the means ± SEM of triplicate measurements.

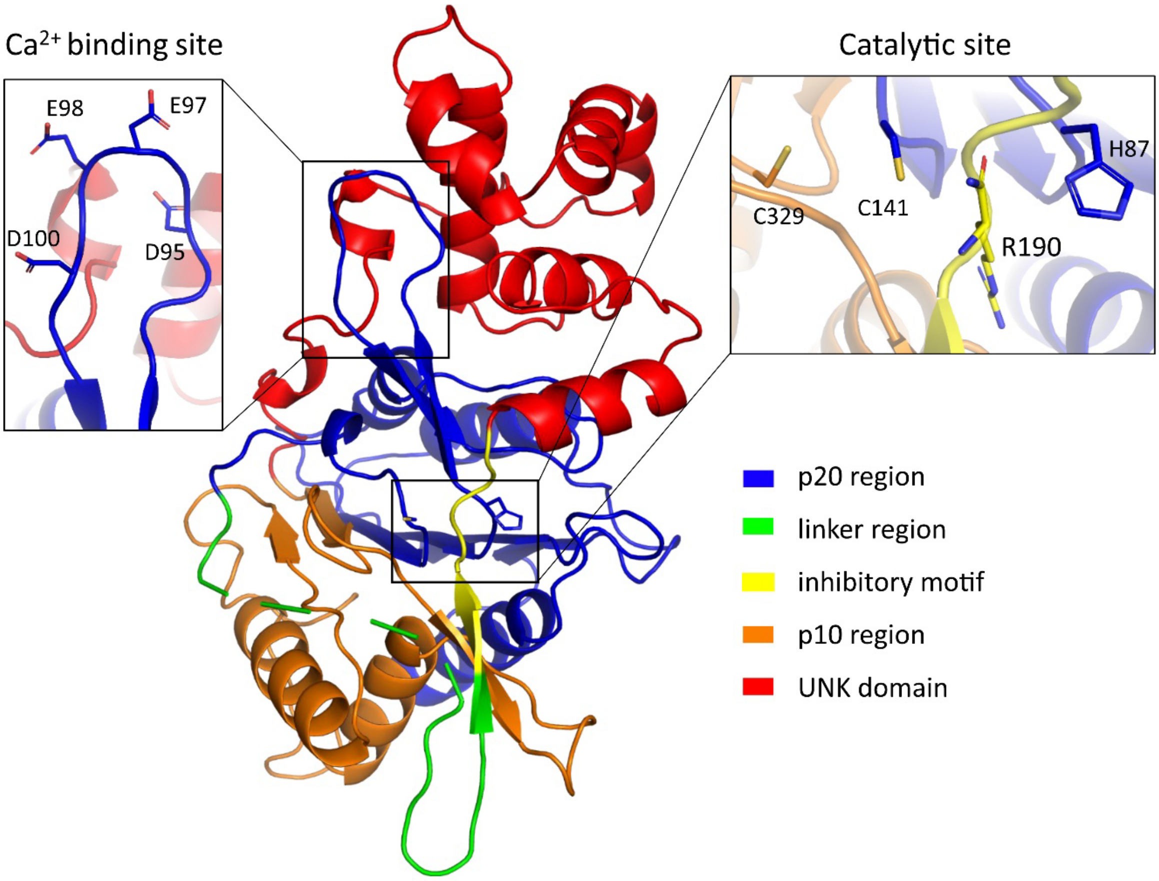


**Supplemental Figure S3** The predicted structure of CrMCA-II. The model was obtained using the Phyre2 web portal (Kelley et al., 2015) in One-to-One threading mode with the AtMCA-IIa structure. UNK (unknown) domain was previously included into the linker region of type II MCAs, but is now defined as a separate structural element (Stael et al., 2023).

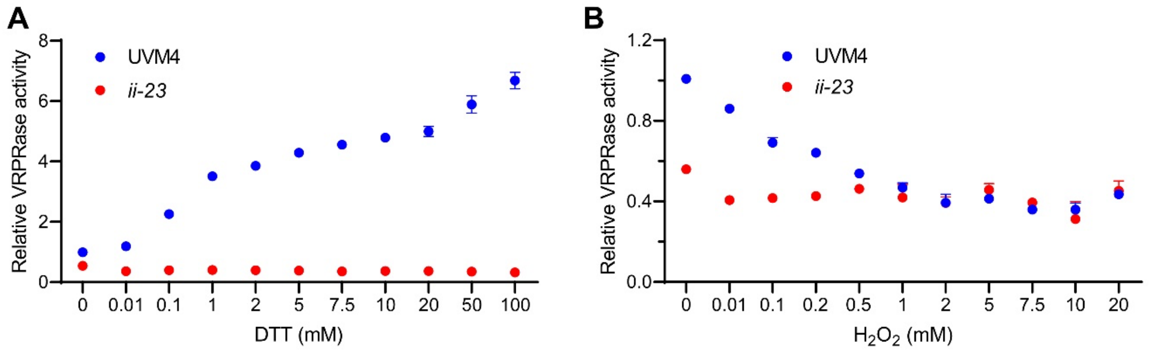


**Supplemental Figure S4** Effect of increased concentration of DTT (**A**) and $H_2O_2$ (**B**) on VRPRase activity in cell lysates prepared from control (UVM4) and CrMCA-II-deficient (*ii-23*) strains of Chlamydomonas. Proteolytic activity was measured under optimal buffer conditions (except varying concentration of DTT in **A**) using Ac-VRPR-AMC as a substrate. Data represent the means ± SEM of triplicate measurements.

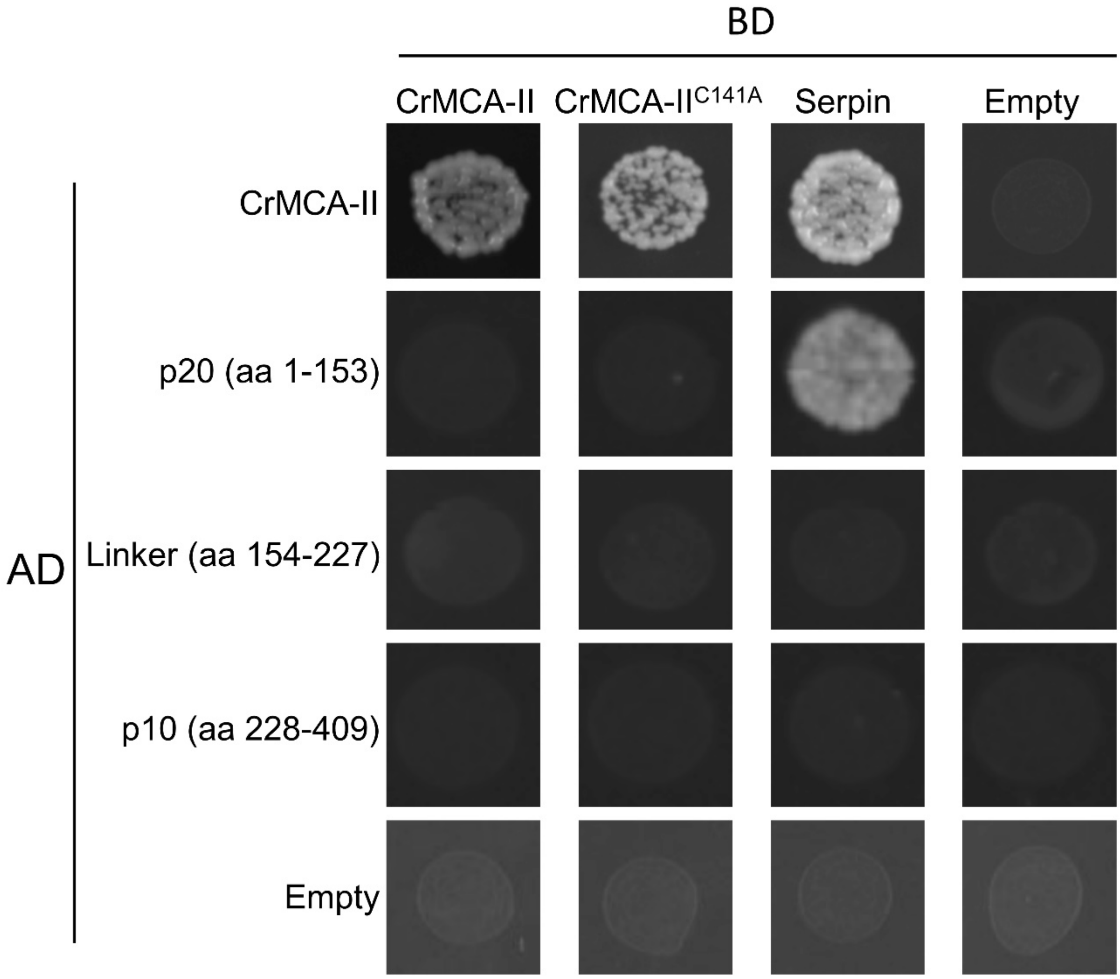


**Supplemental Figure S5** The Y2H assay of interaction between full-length CrMCA-II or its catalytically inactive mutant CrMCA-II$^{C141A}$ and different protein regions. Interaction between full-length CrMCA-II and serpin, an *in vivo* inhibitor of plant MCAs, was used as a positive control. The top row images are from Fig. 1E and are included for comparison.

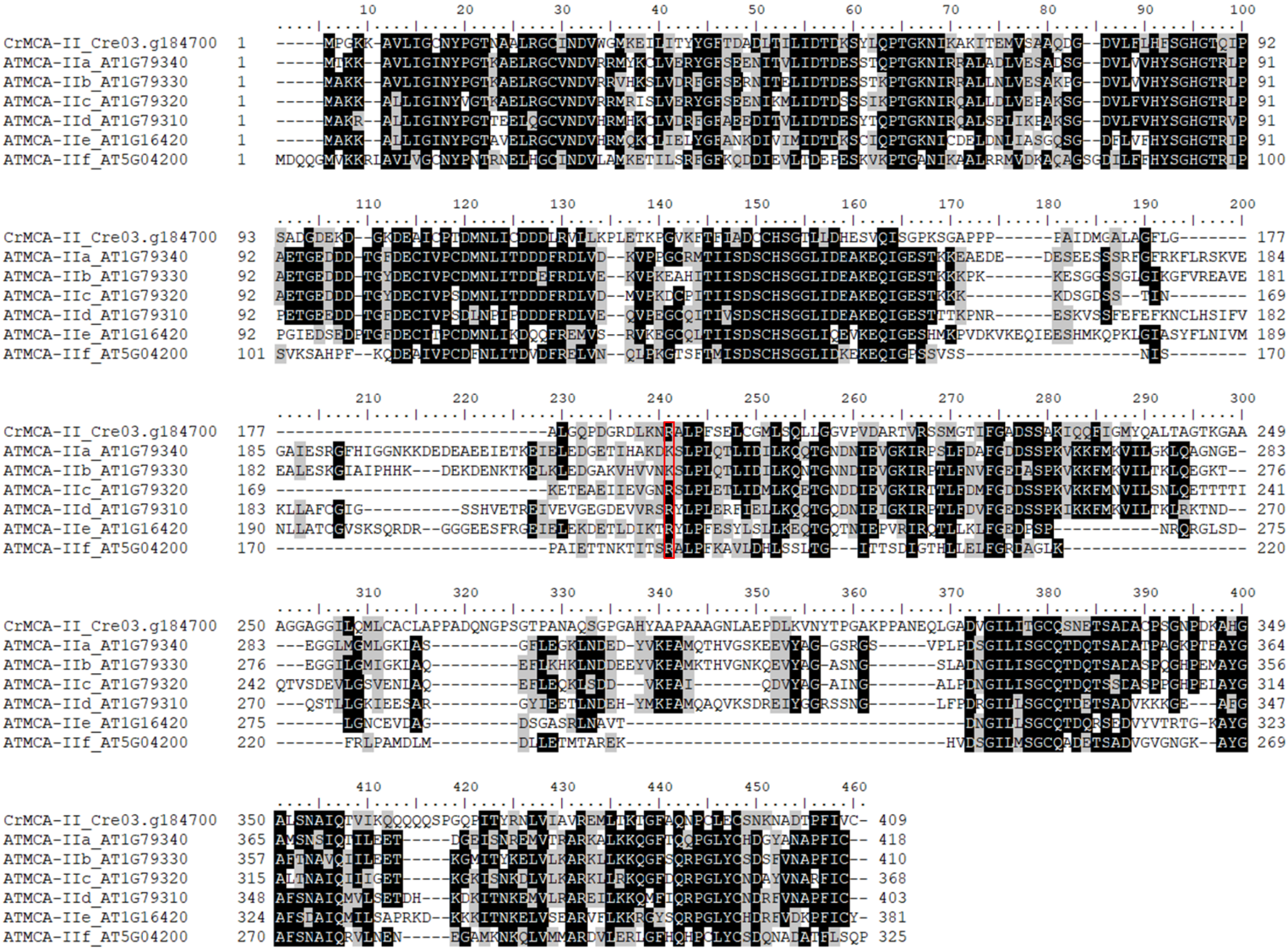


**Supplemental Figure S6** Sequence alignment of type II MCAs from Chlamydomonas and Arabidopsis. The sequences were aligned using ClustalW (Larkin et al., 2007) and visualized by BioEdit (Hall, 1999). Identical amino acid residues are dark shaded and amino acid residues with similar traits are light shaded. Numbers on the left and right sides indicate the position of the first and last amino acid in the full-length protein sequence, respectively. The conserved Arg/Lys auto-cleavage site is denoted by red rectangle.

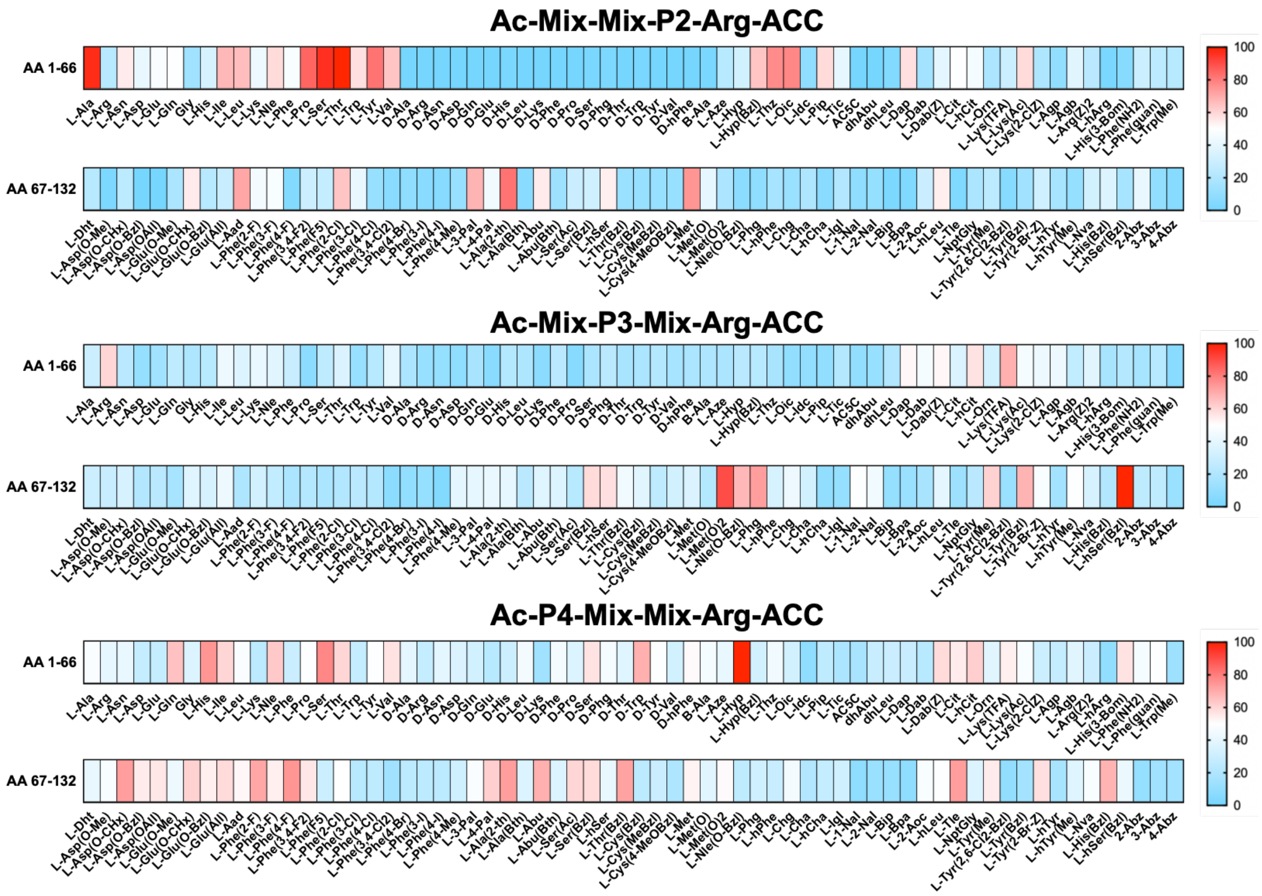


**Supplemental Figure S7** Substrate specificity of rCrMCA-II in the P4-P2 positions. The catalytic preferences of rCrMCA-II were determined using the P1-Arg HyCoSuL library, and are presented as heat-maps. The x-axis shows abbreviated amino acids (from 1 to 132), and the heat-maps display the relative activity of each substrate adjusted to the best-recognized amino acid in each position marked in red. Each sub-library (P2, P3, P4) screening was performed in triplicate under optimal buffer conditions and the rate of substrate hydrolysis are presented as average (S.D. for each measurement was below 10%).

Ac-Val-Asp-Pro-Arg-ACC

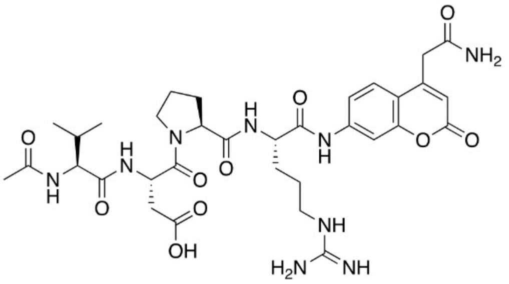

Ac-His-Agp-Met-Arg-ACC

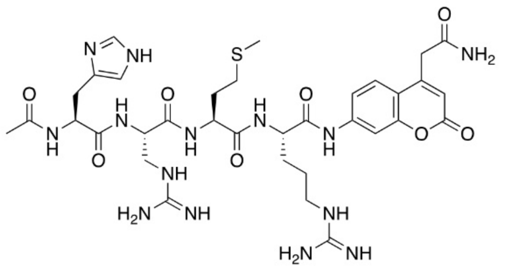

Ac-Val-Arg-Pro-Arg-ACC

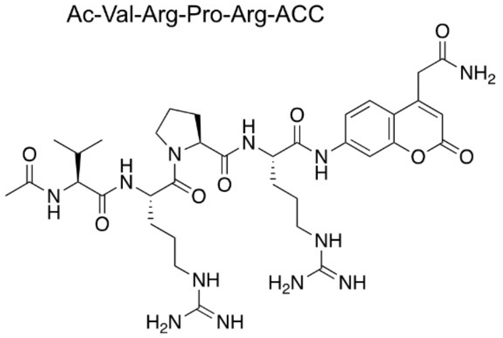

Ac-His(Bzl)-Agp-Ala-Arg-ACC

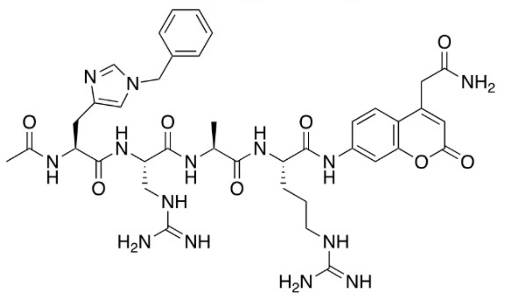

Ac-Ser-Arg-Thr-Arg-ACC

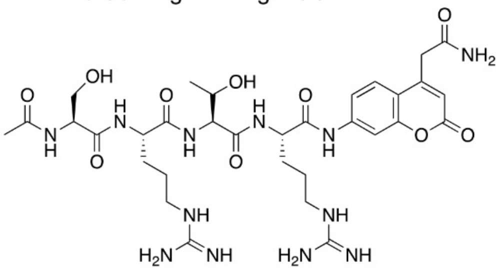

Ac-His-Dap-Ala-Arg-ACC

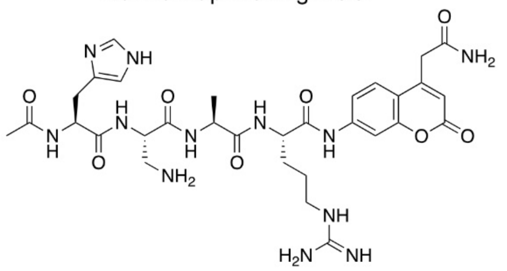

Ac-His-Arg-Thr-Arg-ACC

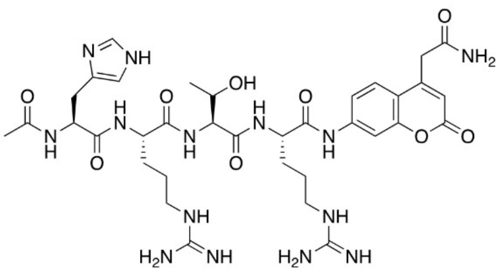

Ac-His(Bzl)-Arg-Thr-Arg-ACC

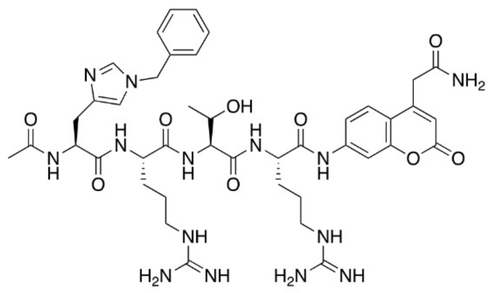

Ac-hP-hS(Bzl)-Thr-Arg-ACC

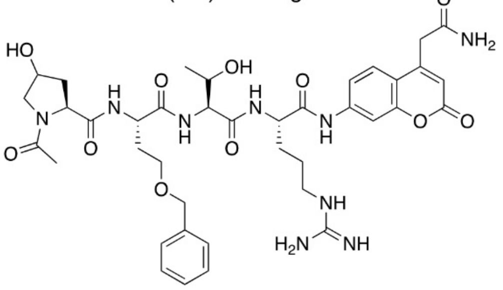

Ac-Ser-Agp-Thr-Arg-ACC

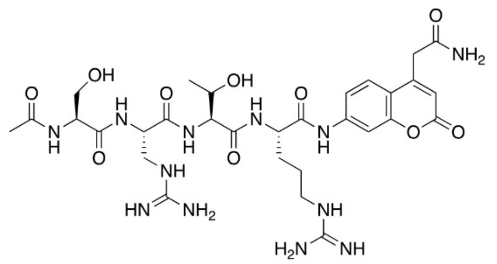

Ac-Ser-Met(O$_2$)-Thr-Arg-ACC

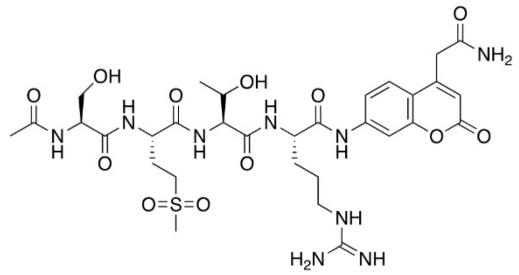

Ac-His(Bzl)-hSer(Bzl)-Thr-Arg-ACC

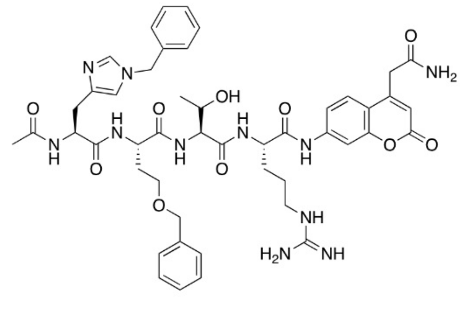

Ac-Ser-Tyr(Bzl)-Thr-Arg-ACC

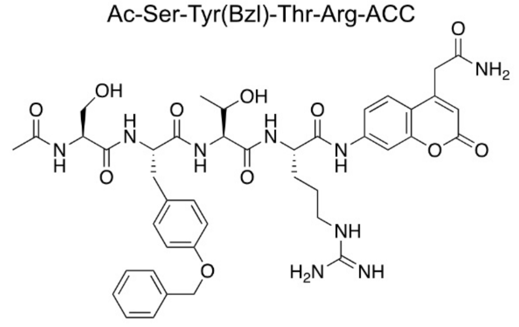

Ac-His-hSer(Bzl)-Met-Arg-ACC

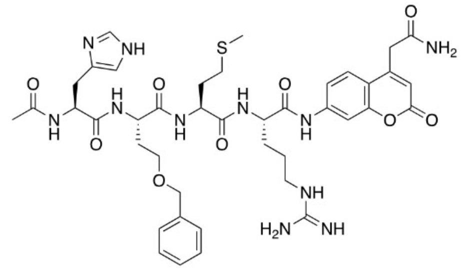

Ac-Ser-hS(Bzl)-Pro-Arg-ACC

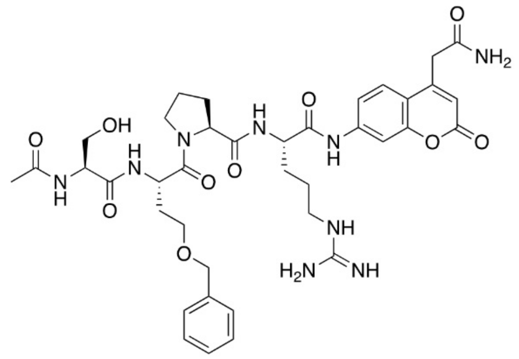

Ac-Ser-hSer(Bzl)-Thr-Arg-ACC

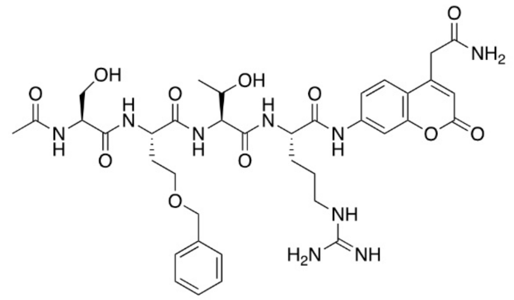

Ac-His(Bzl)-Dap-Met-Arg-ACC

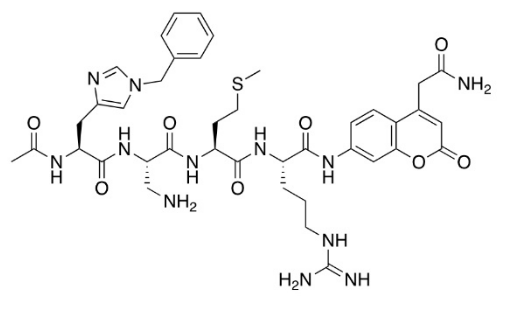

**Supplemental Figure S8** Structures of ACC-labeled tetrapeptide substrates for CrMCA-II.

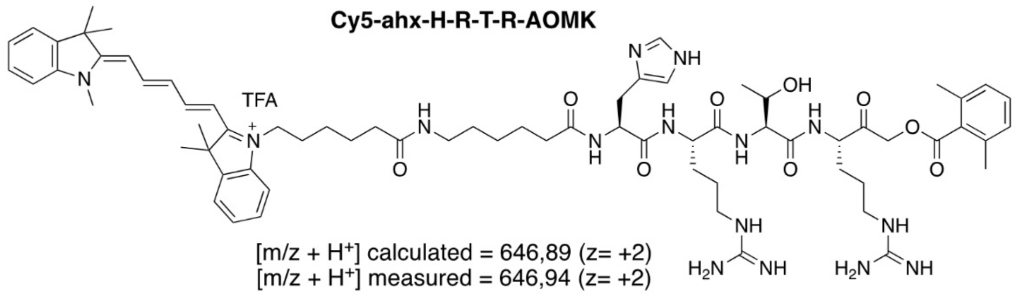


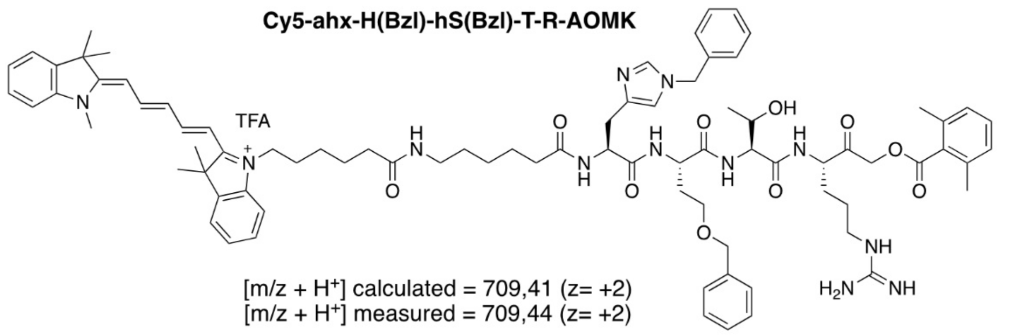


**Supplemental Figure S9** Structures and MS data of Cy5-labeled ABPs for CrMCA-II.

```
UVM4    GCCAAGATCACGGAGATGGTGAGCGCCGCCCAGGACGGGGATGTGCTGTTCCTGCACTTCAGCGGCCACGGAACCCAGATCCCCAGCGCAGACGGCGACGAGAAGG
ii-9    GCCAAGATCACGGAGATGGTGAGCGCCGCCCAGGACGGGGATGTGCTGTTCC-paro cassette--CGGAACCCAGATCCCCAGCGCAGACGGCGACGAGAAGG
ii-23   GCCAAGATCACGGAGATGGTGAGCGCCGCCCAGGAC---------paro cassette----------CGGAACCCAGATCCCCAGCGCAGACGGCGACGAGAAGG
```

GTGCTGTTCCTGCACTTCAG: gRNA targeting site
CGG: protospacer adjacent motif (PAM)
paro cassette: PSAD promoter + *AphVIII* gene + PSAD terminator(~1370 bp)

**Supplemental Figure S10** The sequence alignment of a part of exon 4 of *CrMCA-II* genomic DNA from control (UVM4) and mutant (*ii-9* and *ii-23*) strains. Paro cassette, paromomycin resistance cassette.

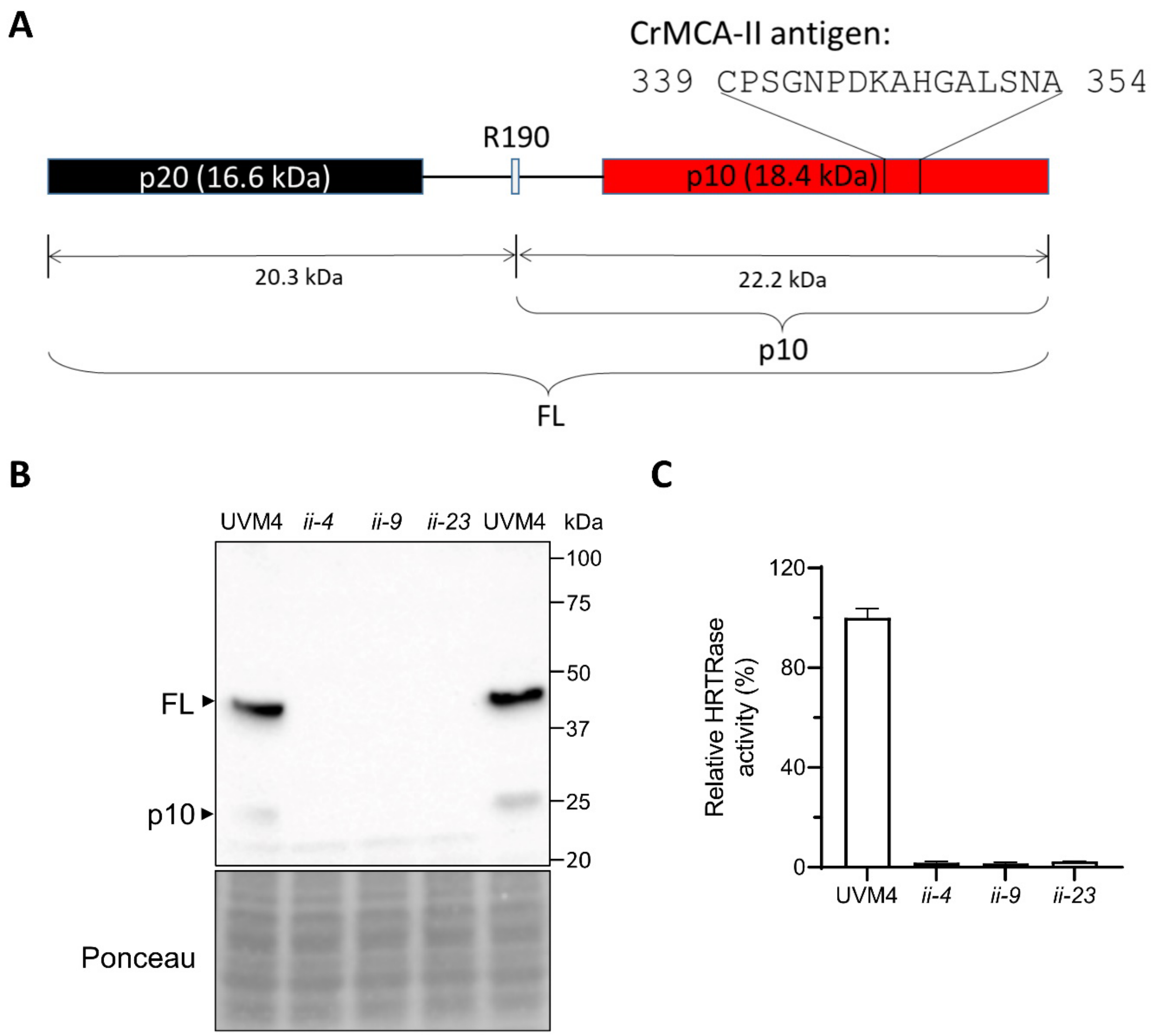


**Supplemental Figure S11** Characterization of *CrMCA-II* mutants using specific antibody and substrate cleavage assay. **A.** Schematic illustration of CrMCA-II domain organization showing location and sequence of peptide used as antigen for generating **α**-CrMCA-II. While p10 region of CrMCA-II is 18.4 kDa, the lower band detected on immunoblots of total protein extracts with **α**-CrMCA-II corresponds to a 22.2 kDa-fragment that includes the p10 and a part of the linker region generated via the auto-cleavage at Arg190. **B.** Representative immunoblot of total protein extracts from UVM4 and *crmca-ii* mutants with **α**-CrMCA-II recognizing zymogen (full-length, FL) and the 22.2 kDa-fragment (indicated as p10). Ponceau staining was used as a loading control. **C.** Ac-HRTR-ACC cleavage assay using cell lysates from UVM4 and *crmca-ii* mutants. Data represent the means ± SEM of triplicate measurements.

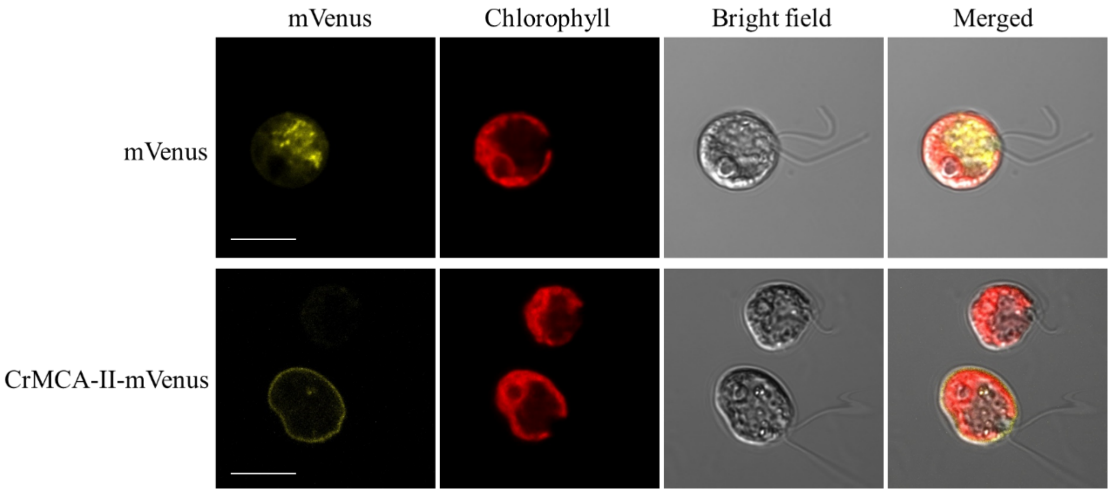


**Supplemental Figure S12** Intracellular localization of mVenus and CrMCA-II-mVenus (yellow) in strain CC-4533 under normal conditions. Scale bars, 10 μm.

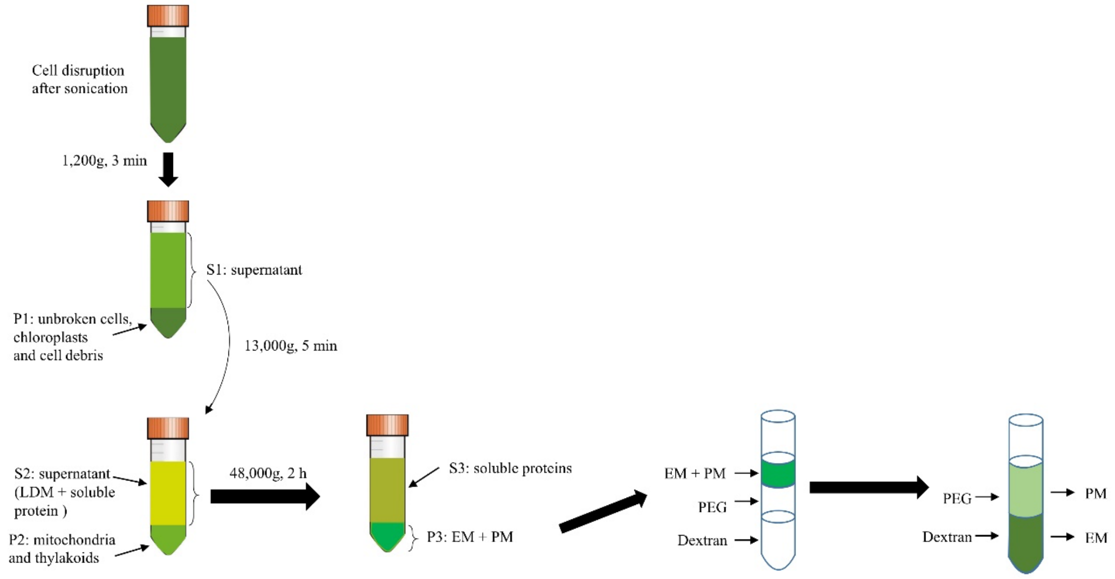


**Supplemental Figure S13** The procedure for subcellular fractionation used for CrMCA-II and $H^+$ATPase immunoblot analysis shown in Figure 5B. EM, endomembrane fraction; PM, plasma membrane fraction. LDM, low-density membranes. PEG, polyethylene glycol.

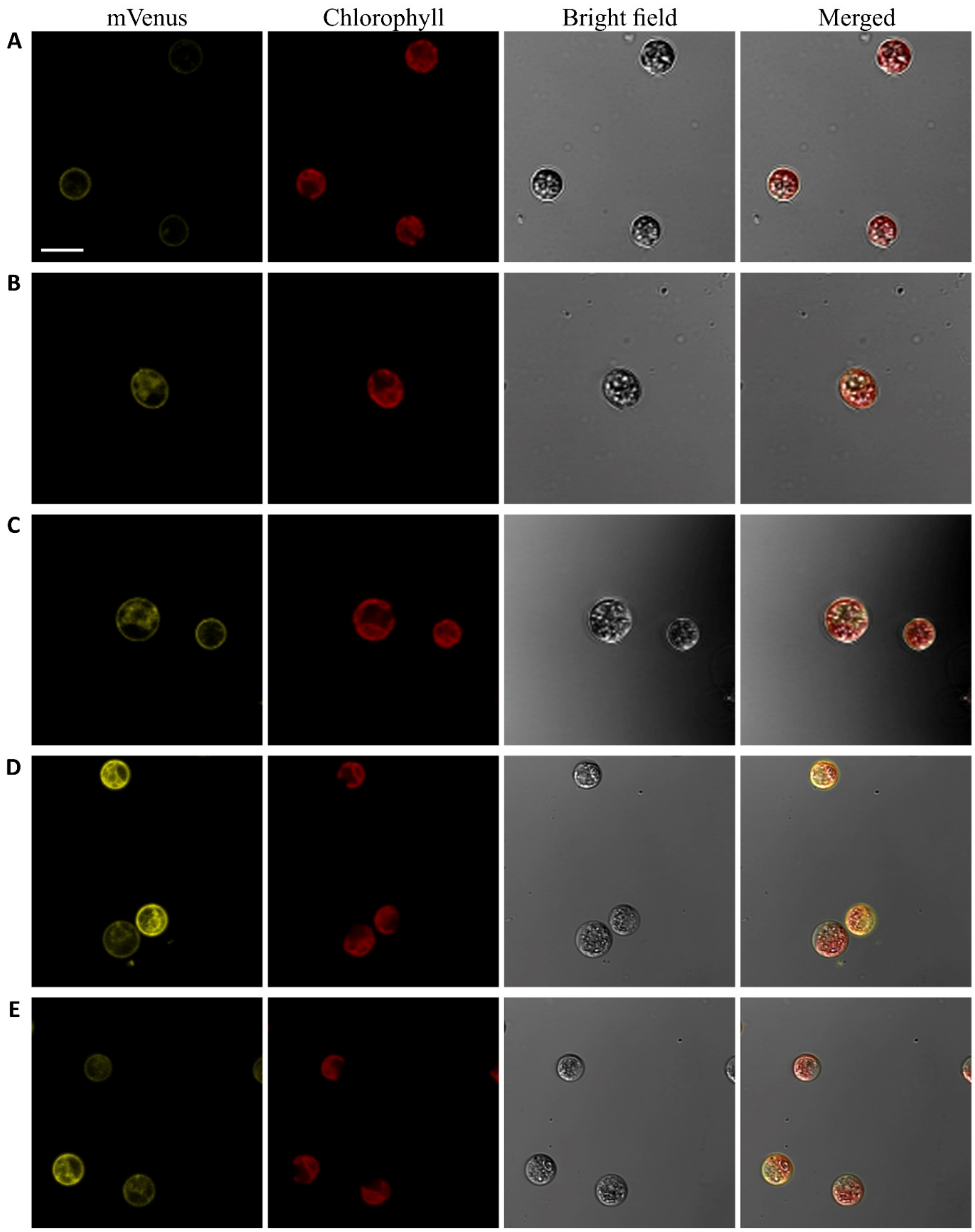


**Supplemental Figure S14** Intracellular localization of CrMCA-II$^{C141A}$-mVenus (yellow) in strain UVM4 before (**A**) and after HS at 42°C for 60 min (**B, C**). **D, E** Localization of CrMCA-II-mVenus under HS in the cells treated with 35 µM cycloheximide. Scale bars, 10 µm.

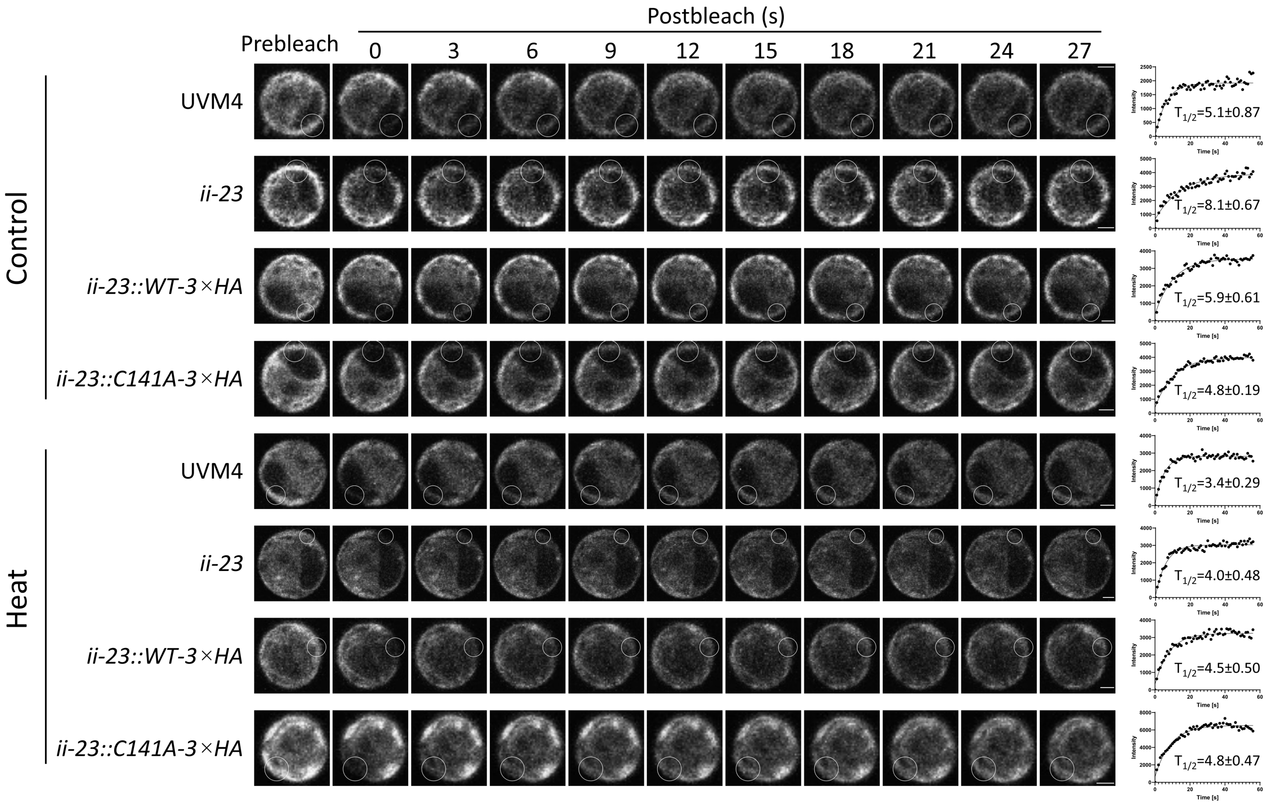


**Supplemental Figure S15** Selected time frames from FRAP analysis shown in Fig. 5F. The rings indicate the bleached foci. Scale bars, 2 µm. Shown on the right are FRAP curves and signal recovery values ($T_{1/2}$; means ± SEM). The signal intensity at postbleach point 0 was set to zero.

**Supplemental Table S1** Mass spectrometry analysis of ACC-labeled tetrapeptide substrates for CrMCA-II. All substrates were purified on semi-preparative HPLC (C8 column) to at least 95% of purity.

| Substrate | $[m/z + H]^+$ calculated | $[m/z + H]^+$ measured |
|---|---|---|
| Ac-Val-Asp-Pro-Arg-ACC | 728.34 | 728.38 |
| Ac-Val-Arg-Pro-Arg-ACC | 385.21 (z = +2) | 385.27 (z = +2) |
| Ac-Ser-Arg-Thr-Arg-ACC | 381.19 (z = +2) | 381.11 (z = +2) |
| Ac-His-Arg-Thr-Arg-ACC | 271.14 (z = +3) | 271.18 (z = +3) |
| Ac-hPro-hSer(Bzl)-Thr-Arg-ACC | 822.38 | 822.45 |
| Ac-Ser-Met(O2)-Thr-Arg-ACC | 768.30 | 768.39 |
| Ac-Ser-Tyr(Bzl)-Thr-Arg-ACC | 858.38 | 858.37 |
| Ac-Ser-hSer(Bzl)-Pro-Arg-ACC | 792.37 | 792.32 |
| Ac-His(Bzl)-Dap-Met-Arg--ACC | 431.19 (z = +2) | 431.12 (z = +2) |
| Ac-His-Agp-Met-Arg-ACC | 271.79 (z = +3) | 271.85 (z = +3) |
| Ac-His-Dap-Ala-Arg-ACC | 237.78 (z = +2) | 237.79 (z = +2) |
| Ac-His(Bzl)-Agp-Ala-Arg-ACC | 422.20 (z = +2) | 422.20 (z = +2) |
| Ac-His(Bzl)-Arg-Thr-Arg-ACC | 451.22 (z = +2) | 451.25 (z = +2) |
| Ac-Ser-Agp-Thr-Arg-ACC | 367.17 (z = +2) | 367.23 (z = +2) |
| Ac-His(Bzl)-hSer(Bzl)-Thr-Arg-ACC | 936.44 | 936.49 |
| Ac-His-hSer(Bzl)-Met-Arg-ACC | 438.69 (z = +2) | 438.75 (z = +2) |
| Ac-Ser-hSer(Bzl)-Thr-Arg-ACC | 796.36 | 796.41 |
| Ac-His-hSer(Bzl)-Thr-Arg-ACC | 423.70 (z = +2) | 423.72 (z = +2) |
| Ac-hPro-hSer(Bzl)-Ser-Arg-ACC | 808.36 | 808.39 |

**Supplemental Table S2** The primers used in this study.

| Name | Sequence 5'-3' | Application |
|---|---|---|
| CrMCA-II-F1 | ttgaagacaaaATGCCGGGCAAGAAGG | CrMCA-II genomic DNA cloning |
| CrMCA-II-R1 | ttgaagacaacgaaCACACAATGAACGGGG | |
| CrMCA-II-F2 | ttgaagacaaGAAcACACATGGATCCCG | |
| CrMCA-II-R2 | ttgaagacaagTTCCATAAAATAAGAGCTTAGG | |
| CrMCA-II-F3 | ttgaagacaaGGTtTCCAGCGGCTTCA | |
| CrMCA-II-R3 | ttgaagacaaaACCAAGCCCGGTGTG | |
| CrMCA-II-F4 | TCAACGATGTGTGGGGAATG | qPCR detection of *CrMCA-II* |
| CrMCA-II-R4 | TTGGCCTTGATGTTCTTCCC | |
| RACK1-F | CTTCTCGCCCATGACCAC | qPCR detection of *RACK1* |
| RACK1-R | CCCACCAGGTTGTTCTTCAG | |
| AphVIII-F | GTATCCCGGTTGTGAGTGGG | qPCR detection of *AphVIII* casette |
| AphVIII-R | CCCTCCACAACACGAGGTAC | |
| CrMCA-II crRNA | /AltR1/rGrUrGrCrUrGrUrUrCrCrUrGrCrArCrUrUrCr ArGrGrUrUrUrUrArGrArGrCrUrArUrGrCrU/AltR2/ | crRNA for *CrMCA-II* gene |
| CrMCA-II long arm paromomycin F | AACAGGCCAAGATCACGGAGATGGTGAGCG CCGCCCAGGActgtgggtctcatgccgaat | for cloning AphVIII gene with 40 bp sequences flanking the CrMCA-II editing site |
| CrMCA-II long arm paromomycin R | ACCCTTCTCGTCGCCGTCTGCGCTGGGGATC TGGGTTCCGagcttgaattcttgcagcgc | |
| CrMCA-II-F5 | TTGGAGTTCATGTTATCTTGCCT | colony PCR for detecting the insertion after genome editing |
| CrMCA-II-R5 | ACCGATACTTACATACCCGCAC | |
| Serpin-F | CCGGAATTCATGCCGCCCGTCAAGG | cloning Serpin into BD vector |
| Serpin-R | CGCGGATCCTTCAATCTTGTGAACGACG | |
| CrMCA-II-F6 | atggaggccGAATTCATGCCGGGCAAGAAGG | cloning CrMCA-II into AD and BD vector |
| CrMCA-II-R6 | caggtcgacggatccGCACACAATGAACGGGG | |
| CrMCA-II-C141A-F | tcccagagtgAGCgcagtcggcga | cloning CrMCA-II-C141A into BD vector and MoClo system |
| CrMCA-II-C141A-R | tcgccgactgcGCTcactctggga | |
